# Production and Purification of Endogenously Modified tRNA-Derived Small RNAs

**DOI:** 10.1101/2020.01.21.913749

**Authors:** Aleksej Drino, Vera Oberbauer, Conor Troger, Eva Janisiw, Dorothea Anrather, Markus Hartl, Steffen Kaiser, Stefanie Kellner, Matthias R. Schaefer

## Abstract

During particular stress conditions, transfer RNAs (tRNAs) become substrates of stress-induced endonucleases, resulting in the production of distinct tRNA-derived small RNAs (tsRNAs). These small RNAs have been implicated in a wide range of biological processes, but how isoacceptor and even isodecoder-specific tsRNAs act at the molecular level is still poorly understood. Importantly, stress-induced tRNA cleavage affects only a few tRNAs of a given isoacceptor or isodecoder, raising the question as to how such limited molecule numbers could exert measurable biological impact. While the molecular function of individual tsRNAs is likely mediated through association with other molecules, addressing the interactome of specific tsRNAs has only been attempted by using synthetic RNA sequences. Since tRNAs carry post-transcriptional modifications, tsRNAs are likely modified but the extent of their modifications remains largely unknown. Here, we developed a biochemical framework for the production and purification of specific tsRNAs using human cells. Preparative scale purification of tsRNAs from biological sources should facilitate experimentally addressing as to how exactly these small RNAs mediate the multitude of reported molecular functions.

## Introduction

Transfer RNAs (tRNAs) are crucial adaptor molecules for the decoding of messenger RNA (mRNA) during protein synthesis. Besides fulfilling this canonical function, tRNAs are also the source of a heterogeneous class of small RNAs, often called tRNA-derived small RNAs (tsRNAs), which have been the subject of intense scrutiny in recent years. An increasing body of work has assigned functional relevance to various tsRNAs because their detection and occurrence is associated with cellular stress and immune responses, cell proliferation and differentiation^1, 2^, but also ill-understood phenomena such as RNA-based inheritance of extra-chromosomal information between generations^3–6^. tsRNAs have been extensively sequenced resulting in efforts to correctly map and annotate the copy number and sequence identity of these small RNAs^7^. However, since tRNAs carry various post-transcriptional modifications, RNA modification-related biases largely disqualify sequencing-based methods from quantifying tRNA and tsRNA abundance^8–12^ and, therefore, the full extent of the tsRNA pool in a given biological sample remains largely unknown. Importantly, northern blotting on total RNAs indicated that only 0.1-5 % of a given tRNA isoacceptor becomes processed into tsRNAs^13^, raising the question as to how such low small RNA quantities could be biologically effective in various and seemingly diverse biological processes.

The biogenesis of different tsRNA species can be attributed to endonucleolytic activities targeting pre-tRNAs or matured tRNAs in the tRNA loop structures (D-, anticodon-, variable-, and T-loops). The best understood mechanism of tsRNA production is stress-induced tRNA fragmentation by anticodon nucleases (ACNases), which is a conserved hallmark of the eukaryotic stress response. Two eukaryotic ACNase protein families (RNase A and T2) specifically cleave matured and full-length tRNAs in response to stress. Mammalian cells express Angiogenin (ANG)^14–16^, an RNase A-family enzyme that is kept inactive by binding to its inhibitor, RNH1. Upon stress exposure, RNH1 becomes phosphorylated and releases ANG, which trans-locates from the nucleus to the cytoplasm where the enzyme targets single-stranded RNA sequences in tRNAs with a preference for pyrimidine-purine dinucleotides^17^. ANG cleavage of RNA substrates results in 5’ tsRNAs containing a 2’-3’-cyclic phosphate at their 3’-end and 3’ tsRNAs containing a 5’-OH. The reproducible production of distinct stress-induced tsRNAs has been reported after starvation^18^, oxidative stress^13, 19, 20^, nutritional deficiency^21^, hypoxia and hypothermia^22, 23^, heat shock or irradiation^13, 24, 25^. While many tRNAs could be ANG substrates, stress-induced ANG activity only affects a fraction of a particular tRNA isoacceptor and isodecoder pool^10, 26^. How such limitation is achieved remains unclear. Importantly, tRNAs are the most heavily modified RNAs in any cell type^27, 28^. While modifications in the anticodon loop contribute to the optimization of mRNA decoding, modifications that occur outside the anticodon loop (also called core modifications) serve largely structural roles during tRNA processing and maturation^29^ but have also been implicated in modulating access of endonucleases. Since stress-induced tsRNAs are likely derived from modified tRNAs, they likely also carry chemical modifications. However, the modification status of individual tsRNAs has not been determined yet. In addition, tsRNA functionality has largely been addressed after re-introducing mostly synthetic RNA sequences into various biological systems, thereby ignoring the potential impact of RNA modifications on tsRNA-mediated silencing of complementary RNA reporters^30^, on tsRNA-mediated modulation of embryonic stem cells and early mammalian embryonic development^31, 32^, on tsRNA-mediated regulation of heterochromatin^33^, on tsRNA-mediated suppression of retrotransposons^34, 35^, on tsRNA-mediated translational enhancement of specific proteins^36^ or when probing for protein and RNA binders to specific tsRNA sequences^20, 32, 37–40^. Predictably, various RNA modifications can affect the hybridization behavior of RNAs^29, 41^ or their interactions with proteins^42^ and therefore, it would be advisable to utilize modified tsRNAs, rather than synthetic sequences, when addressing and testing their potential for biological impact.

Here, we set out to develop a biochemical framework to purify large amounts of specific tsRNAs, which might allow addressing the biological function of endogenous tsRNAs. As proof of concept, a scalable purification protocol for specific tsRNAs carrying post-transcriptional modifications was established. To this end, two 5’ tsRNA species derived from tRNA-Gly^GCC^ and tRNA-Glu^CUC^, which are dominantly featured in tsRNA-related literature, were purified from human cells using chromatographic and hybridization-based methods. The post-transcriptional modification status of purified tsRNAs was determined by LC-MS/MS. To show the downstream applicability of this approach, purified tsRNAs were used for RNA affinity capture experiments and for the approximation of actual copy numbers of specific and stress-induced 5’ tsRNAs in a human cell line.

## Results

### Inorganic arsenite-induced tRNA fragmentation coincides with increased cell death

5’ tsRNA-Gly^GCC^ and 5’ tsRNA-Glu^CUC^ are two tsRNA species representing tRNA halves that are often detected in biological samples as very abundant, especially during stress conditions. To robustly produce these tsRNAs, HEK293T cells were treated with inorganic sodium arsenite (iAs), whose impact on cellular stress responses can be monitored by the amount of phosphorylated eukaryotic initiation factor 2α (Figure 1A). Northern blotting for tRNA-Gly^GCC^ and tRNA-Glu^CUC^ showed robust tRNA cleavage during the acute stress response to iAs with increased tsRNA levels being detectable 24 hours after removing the stressor (Figure 1B). Of note, dose response measurements taken 24 hours after exposure to acute iAs exposure revealed reduced cell viability exactly at iAs concentrations (between 250-500 μM, supplementary Figures 1A and B) that are commonly used as a conduit for the reproducible induction of stress granule formation and tRNA fragmentation. These results indicated limits to using the iAs-mediated oxidative stress response for producing large numbers of tsRNAs without affecting cell viability.

**Figure 1.**
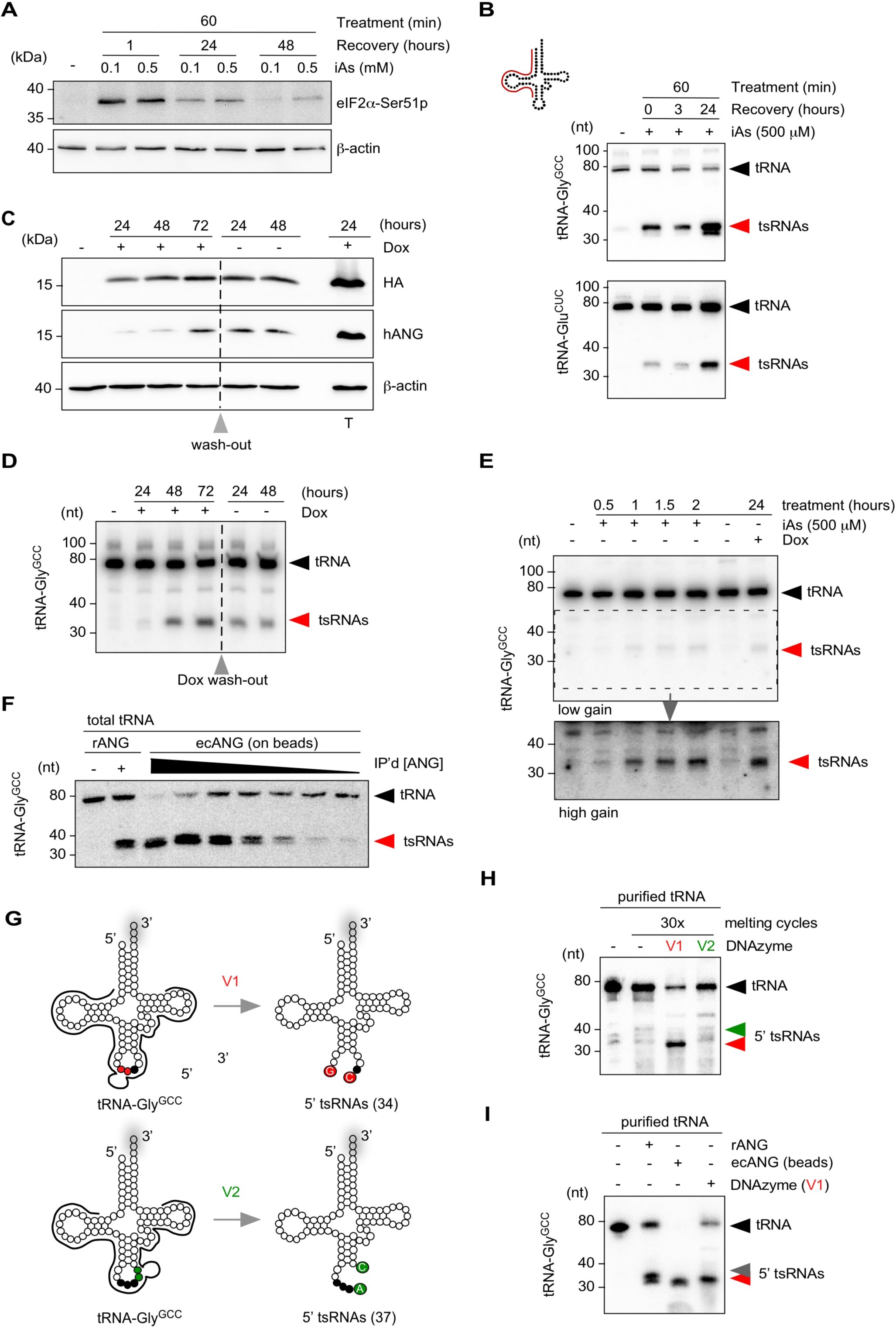
*In vivo* and *in vitro* production of tsRNAs. (A) Western blotting on HEK293T cell protein extract before and after exposure to inorganic arsenite (iAs) using antibodies against phosphorylated eIF2α and β-Actin as a loading control. Cells were treated at 70 % confluency for one hour with 0.1 or 0.5 mM iAs followed by wash-out of iAs and recovery for up to 24 hours. (B) Northern blotting of total RNA (3 µg) extracted from HEK293T cells exposed to 0.5 mM iAs using 5’ probes against tRNA-Gly^GCC^ and tRNA-Glu^CUC^ (annealing of probe in full-length tRNA according to cartoon). Black arrowheads: full-length tRNAs; red arrowheads: tsRNAs. (C) Western blotting on HEK293T-ecANG cell protein extracts obtained from cells that were exposed to Doxycycline (Dox) for three consecutive days before wash-out (grey arrowhead) and further incubation for two days. In addition, HEK293T cells transiently transfected with a Dox-inducible ANG-containing plasmid were induced with Dox (T). Membranes were probed with antibodies against the epitope-tag (HA), against ANG (hANG) and β-Actin as a loading control. (D) Northern blotting of total RNA (3 µg) extracted from HEK293T-ecANG cells exposed to Dox and cultured without Dox (grey arrowhead: wash-out) using a 5’ probe against tRNA-Gly^GCC^ as described in (B). Black arrowheads: full-length tRNAs; red arrowheads: tsRNAs. (E) Northern blotting of total RNA (2.5 µg) extracted from HEK293T cells exposed to iAs for different times and HEK293-ecANG cells exposed to Dox for 24 hours and using a 5’ probe against tRNA-Gly^GCC^ as described in (B). Lower image (high gain) represents a digitally enhanced region of the upper image (low gain) for better visualization of tsRNAs. Black arrowhead: full-length tRNAs; red arrowhead: tsRNAs. (F) Northern blotting of total tRNA fraction (500 ng) subjected to recombinant ANG (100 ng) or ecANG precipitated from HEK293T-ecANG cell culture supernatant. Different amounts of precipitated ecANG on beads were exposed to tRNAs followed by probing against tRNA-Gly^GCC^ as described in (B). Black arrowhead: full-length tRNAs; red arrowhead: tsRNAs. (G) Cartoon summarizing the design of two DNAzymes (10–23 variant), which address the phosphodiester bonds between G34-C35 (v1, red dots) or A37-C38 (v2, green dots) in tRNA-Gly^GCC^, respectively. Black dots depict the anticodon triplet, while grey shadows indicate 3’ CCA-addition in full-length tRNAs. (H) Northern blotting of purified tRNA-Gly^GCC^ (500 ng) subjected to 30x melting and DNAzyme activity cycles followed by probing against tRNA-Gly^GCC^ as described in (B). Black arrowhead: full-length tRNAs; red arrowhead: 5’ tsRNAs resulting from DNAzyme (v1); green arrowhead: position of expected 5’ tsRNAs resulting from DNAzyme (v2) activity. (I) Northern blotting of purified tRNA-Gly^GCC^ (500 ng) subjected to recombinant ANG (100 ng), ecANG precipitated from HEK293T-ecANG cell culture supernatant or DNAzyme activity (v1, 30x melting cycles) using a probe against tRNA-Gly^GCC^ as described in (B). Black arrowhead: full-length tRNAs; red arrowhead: 5’ tsRNAs common to all approaches; grey arrowhead: additional 5’ tsRNA species only detectable using recombinant ANG.

### Overexpression of human ANG results in robust tsRNA production

In an attempt to increase tsRNA production, while avoiding excessive stress-induced cell death and to overcome the negative regulation of endogenous ANG, an inducible ANG-expression system was established. Using the Flp-In™-T-REx™ system, human ANG-HA-FLAG (ecANG) was stably inserted into the genome of HEK293T cells. Addition of doxycycline (Dox) induced robust ecANG expression (Figure 1C) and resulted in the production of 5’ tsRNAs that were comparable in size distribution to iAs-induced 5’ tsRNAs (Figures 1D and E). Importantly, while ecANG localized to visible granules (supplementary Figure 1C), overexpression of ANG did not cause lethality and slowed down cell proliferation in a statistically significant manner only after three days of constant Dox induction (supplementary Figure 1D). These results indicated tolerance of HEK293 cells to massively increased ANG levels but also showed that ectopic ANG was insufficient to fragment a given tRNA isoacceptor to completion, suggesting effective cellular control mechanisms which limit excessive ANG activity.

### Production of tsRNA using secreted ecANG

Recombinant ANG has been used to fragment purified tRNA *in vitro*^22, 43^. Since ANG is secreted from cells^44, 45^, cell culture supernatants from HEK293 cells expressing ecANG were collected and ecANG was immuno-precipitated *via* its FLAG-tag (supplementary Figure 1E). Precipitated ecANG was used for tsRNA production on gel-purified total tRNAs. Northern blotting on the cleavage reactions using probes against the 5’ half of tRNA-Gly^GCC^ showed that secreted and immuno-precipitated human Angiogenin can be used to produce scalable quantities of specific tsRNAs *in vitro* (Figure 1F and supplementary Figure 1F).

### Production of specific tsRNA using DNAzymes

DNAzymes are short deoxyribonucleic acids displaying RNA hydrolyzing activity, which can be designed to cleave RNAs with site-directed specificity. As an alternative to ANG-mediated cleavage of tRNAs, DNAzymes of the 10–23 variant^46–48^ were designed to target human tRNA-Gly^GCC^ between G34-C35 (variant 1, v1) or A37-C38 (variant 2, v2), respectively (Figure 1G). Small RNAs were purified from HEK293 cells using ion-exchange chromatography (anIEX) followed by RNA affinity capture using 5’ covalently immobilized amino-modified DNA oligonucleotides complementary to the 5’ end of tRNA-Gly^GCC^. Purified tRNA-Gly^GCC^ was subjected to DNAzyme activity *in vitro*. Northern blotting using probes against the 5’ half of tRNA-Gly^GCC^ showed robust and almost quantitative hydrolysis into tsRNA when using v1 (Figure 1H). In contrast, v2 did not hydrolyze tRNA-Gly^GCC^ to an appreciable extent indicating that specifically tailored DNAzymes can be used for the production of specific tsRNAs *in vitro*. Of note, previous attempts targeting the phosphodiester bond between positions 37-38 in a different tRNA (*Drosophila melanogaster* tRNA-Asp^GUC^) yielded also limited cleavage^48^ indicating a common interference with DNAzyme efficiency at exactly this position. While the purine-pyrimidine context at this position in both human tRNA-Gly^GCC^ and *Drosophila* tRNA-Asp^GUC^ is ApC, this dinucleotide is the least addressable context for DNAzymes according to^46^. In addition, C38 in both tRNAs is modified by Dnmt2/TRDMT1 proteins, which might negatively affect cleavage yield as has been observed for other RNA modifications^49^. When comparing the efficiencies of tRNA fragmentation mediated by recombinant ANG-, immuno-purified ecANG- or DNAzyme, immuno-purified ecANG was able to quantitatively fragment tRNA-Gly^GCC^ (Figure 1I). Of note, while the DNAzyme produced a detectable 5’ tsRNA of 34 nucleotides, targeting recombinant ANG and immunprecipitated ecANG to tRNA-Gly^GCC^ *in vitro* resulted in slightly different 5’ tsRNAs (Figure 1I). ANG hydrolyses preferentially CpA in single-stranded RNA, but also CpC and CpG dinucleotides can be targeted^17, 50, 51^. Human tRNA-Gly^GCC^ contains C35-C36, C36-A37 and a C38-G39 in the anticodon and ANG-mediated creation of two closely migrating 5’ tsRNAs could be the result of structural changes caused by *in vitro* melting and refolding of tRNAs or by differences in the activities of recANG and ecANG. In addition, ANG cleavage creates 5’ tsRNAs containing a 2’-3’-cyclic phosphate (cP) at their 3’ ends, which increases their migration behavior. Importantly, two distinct 5’ tsRNA species can sometimes be observed also *in vivo*, especially during the stress recovery or when exposing cells to exceedingly high iAs concentrations (Figure 1, supplementary Figure 1B and Figures below), which could cause stochastic cP ring opening since cP not being very stable in aqueous solutions and susceptible to background hydrolysis^52^.

### Purification of specific stress-induced and ectopic ANG-produced tsRNAs

To produce tsRNAs in scalable quantities from endogenous sources, 5’ tsRNA-Gly^GCC^ and 5’ tsRNA-Glu^CUC^ were purified from iAs-treated or ecANG-expressing HEK293 cells by anIEX and affinity capture using 5’ amino-modified DNA oligonucleotides (complementary to the target RNA) covalently immobilized on NHS-linked sepharose columns (Figure 2A). Denaturing PAGE and northern blotting for target tRNAs (i.e., tRNA-Gly^GCC^) showed clear enrichment of 5’ tsRNAs after the affinity capture step (Figure 2B). To separate 5’ tsRNAs from residual co-purified parental tRNAs, preparative SEC (prepSEC), RNaseH-mediated RNA removal or urea-PAGE gel elution was used (Figures 2C and D). Both prepSEC and gel elution followed by RNA precipitation resulted in very reproducible tsRNA content. The calculated yield of such an RNA purification indicated that about 1-2 μg of a particular 5’ tsRNA species (about 90-180 pmoles) can be purified from the small RNA fraction obtained from about 100 million HEK293T cells (ca. 200 μg total RNA) after exposure to iAs or expression of ecANG.

**Figure 2.**
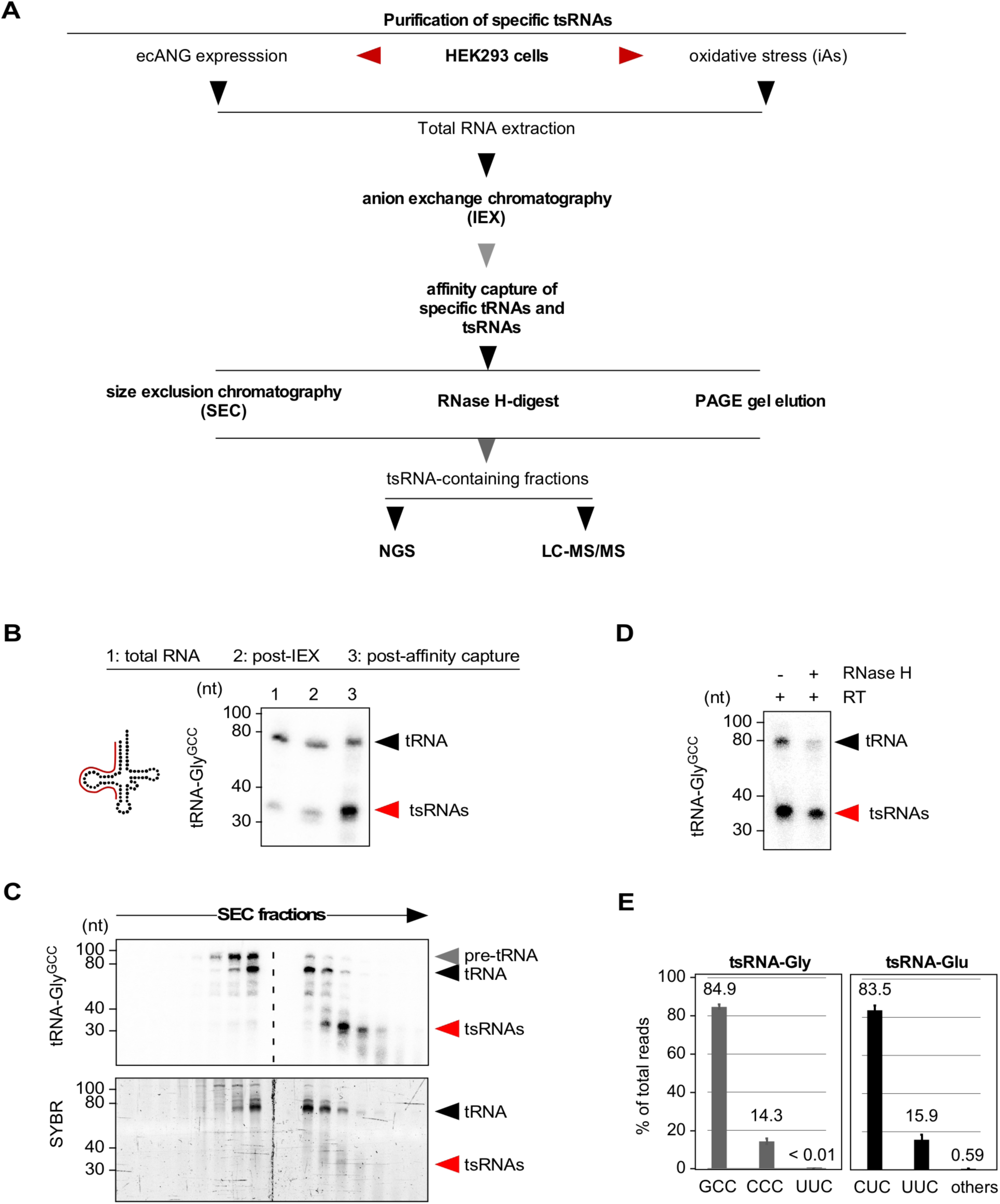
Purification of specific tsRNAs from HEK293 cells. (A) Flowchart summarizing the purification of specific tsRNAs from HEK293T cells exposed to acute stress conditions (iAs) or HEK293T cells expressing ecANG for 48 hours. (B) Northern blotting of total RNA (3 µg) after small RNA enrichment using anion-exchange chromatography (post-IEX) and affinity capture of tRNA-Gly^GCC^-derived sequences from the small RNA pool using a 5’ probe against tRNA-Gly^GCC^ (annealing of probe in full-length tRNA according to cartoon). Black arrowhead: full-length tRNAs; red arrowhead: tsRNAs. (C) Northern blotting of RNA fractions obtained from size exclusion chromatography after affinity capture of tRNA-Gly^GCC^-derived sequences as described in (A) using a 5’ probe against tRNA-Gly^GCC^. Lower panel shows the SYBR-stained PAGE-resolved RNA fractions containing both tRNAs and tsRNAs. Black arrowheads: full-length tRNAs; red arrowhead: tsRNAs. (D) Northern blotting of RNAs using a 5’ probe against tRNA-Gly^GCC^ after affinity capture of tRNA-Gly^GCC^-derived sequences as described in (A), reverse transcription (RT) with a primer annealing to the 3’-end of tRNA-Gly^GCC^ followed by RNase H-digestion of full-length but not 5’ tsRNAs. Black arrowhead: full-length tRNAs; red arrowhead: tsRNAs. (E) Bar-chart depicting the distribution of all small RNA sequencing reads (in %) from affinity-captured and SEC-purified tRNA-Gly^GCC^-and tRNA-Glu^CUC^-derived sequences.

### Determining the identity of purified 5’ tsRNAs

Northern blotting of purified 5’ tsRNA-Gly^GCC^ for other tRNA sequences showed very low cross-reactivity with 3’ tsRNA-Gly^GCC^ or 5’ tsRNA-Glu^CUC^ sequences indicating low contamination of the affinity capture eluate with other tRNA sequences (supplementary Figure 2A). Triplicate small RNA sequencing of ecANG-produced 5’ tsRNA-Gly^GCC^ revealed that 84.9 % of all reads mapped to 5’ tsRNA-Gly^GCC^ while 14.3 % were derived from the 5’ end of the isoacceptor tRNA-Gly^CCC^ (particularly from tRNA-Gly^CCC-1.1^ and tRNA-Gly^CCC-1.2^, which share 100 % sequence identity with the 5’ half of tRNA-Gly^GCC^, Figure 2E and supplementary Figure 2B). Similarly, after purification of 5’ tsRNA-Glu^CUC^ from ecANG cells, 83.5 % of all reads mapped to 5’ tsRNA-Glu^CUC^ while 15.9 % were derived from the 5’ end of the isoacceptor tRNA-Glu^UUC^, which shares 96 % sequence identity with the 5’ half of tRNA-Glu^CUC^ (Figure 2E and supplementary Figure 2B). These results indicated that purification of particular 5’ tsRNAs can be achieved with only minor contamination from other RNAs including other tRNA-derived sequences although we cannot absolutely exclude the co-purification of additional RNAs since sequencing-based RNA identification approaches suffer from amplification biases caused by particular RNA modifications (such as m^1^A), which often interfere with reverse transcription^12^ thereby leading to an underrepresentation of particular sequencing reads and therefore an underestimation of RNA existence in the biological sample. Since the hybridization-based purification of particular tRNA-Gly and tRNA-Glu isoacceptors produced about 15 % co-purified tRNA isoacceptor sequences (Gly^CCC^ and Glu^UUC^, respectively), we will, from here on, label tRNA-derived sequences as tRNA or 5’ tsRNA for Gly^GCC/CCC^ and Glu^CUC/UUC^.

### Determining the modification status of purified tsRNAs

LC-MS/MS analyses can reproducibly quantify individual RNA modifications in purified tRNAs as shown by analyzing six LC-MS/MS measurements on commercially available tRNA-Phe from *Saccharomyces cerevisiae* (supplementary Figures 3A and B confirming the analysis of *S.c.* tRNA-Phe published in^53^). 5’ tsRNA-Gly^GCC/CCC^ and 5’ tsRNA-Glu^CUC/UUC^ along with their remaining parental full-length tRNAs were purified from biological triplicate experiments using either iAs exposure or ecANG expression followed by LC-MS/MS analyses (Figure 3, supplementary Tables 1 and 2). Absolute quantification of modifications in full-length tRNAs and 5’ tsRNAs revealed reproducible modification levels of modifications predicted to reside in the 5’ halves of parental tRNAs (N2’-methylguanosine, m^2^G in both 5’ tsRNAs and 2’-O-methyluridine, Um in 5’ tsRNA-Gly^GCC/CCC^)^27^ albeit with low stoichiometry suggesting incompletely modified tRNA and 5’ tsRNA molecules. Importantly, RNA modifications not expected in purified tRNAs and tsRNAs (i.e., m^7^G, m^1^G, m^22^G, Am, m^6^A) were either not present or detected at very low levels (Figures 3C-G, supplementary Table 2), supporting the notion that affinity capture was enriching for targeted RNAs. In addition, tRNA modification levels predicted to reside in the 3’ halves of full-length tRNAs (3’ of the ANG cleavage sites in the anticodon loop) such as m^5^C, m^1^A or m^5^U were very low to non-existent in purified tsRNAs (Figures 3C-G, supplementary Table 2). Of note, quantification of dihydrouridine (D) was not possible due to presence of D in the digestion cocktail introduced through the deaminase inhibitor tetrahydrouridine. Interestingly, levels of pseudouridine (Y) in both 5’ tsRNAs were consistently lower than in co-purified parental tRNAs. These combined results indicated that ANG activity induced through iAs-exposure or by ectopic expression was largely directed towards tRNAs containing particular modification patterns such as low Y content. Furthermore, these results showed that no additional RNA modifications, which were not present in parental tRNAs, were added to 5’ tsRNAs in the process of tRNA cleavage.

**Figure 3.**
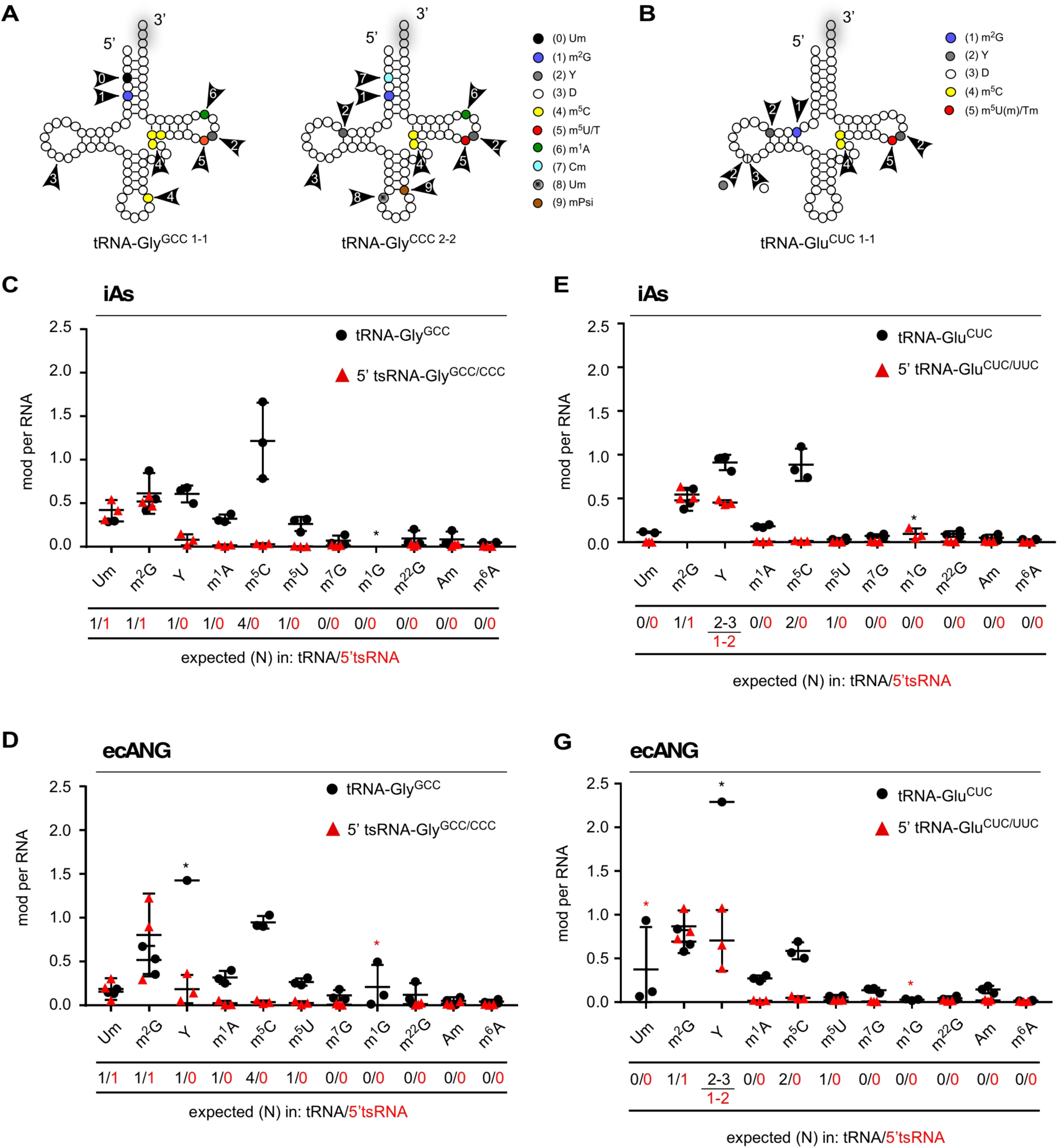
Determination of RNA modification patterns in purified tsRNAs. (A-B) Cartoon depicting known RNA modifications in tRNA-Gly^GCC-1.1^/tRNA-Gly^CCC-2.2^ and tRNA-Glu^CUC-1.1^ (according to http://gtrnadb.ucsc.edu/), which can be captured by NHS-coupled complementary oligonucleotides (see supplementary Figure 2). Arrows and numbers show color-coded modifications at specific positions. (C-D) Results from triplicate LC-MS/MS analysis of tRNA-Gly^GCC/CCC^-derived sequences purified from HEK293T cells that were exposed to acute stress conditions (iAs) or HEK293T-ecANG cells that were induced with Dox for 48 hours before RNA extraction and affinity capture of tRNA-derived sequences. tsRNAs and remaining parental tRNAs were co-purified from the same small RNA pool. (E-F) Results from triplicate LC-MS/MS of tRNA-Glu^CUC/UUC^-derived sequences purified from HEK293T cells that were exposed to acute stress conditions (iAs) or HEK293T-ecANG cells that were induced with Dox for 48 hours before processing as in (C-D). Tables below charts depict expected and detected levels of RNA modifications per tRNA and tsRNA according to^27^. From three replicate measurements, plotted as single data points with mean and standard deviation. Asterisk denotes that signal in 1-3 replicates was below lower limit of quantification.

### Identification of proteins associating with endogenous 5’ tsRNAs

RNA affinity capture of interacting proteins from cell extracts was performed using 5’ tsRNA-Glu^CUCUUC^ purified from HEK293T-ecANG cells. To this end, 5’ tsRNA-Glu^CUC/UUC^ was 5’ biotinylated and coupled to streptavidin-coated sepharose beads. Since virtually all iAs-induced 5’ tsRNA-Gly^GCC/CCC^ and 5’ tsRNA-Glu^CUC/UUC^ resided in the cytoplasm (Figures 4A and B), fractionated cytoplasmic protein extracts (CPEs) from HEK293T cells growing under steady-state conditions or from cells subjected to acute iAs exposure were used for RNA affinity capture experiments. Mass spectrometry analysis on tryptic digests and label-free quantification (LFQ) of peptides was performed against empty streptavidin matrix (noRNA control) and an unrelated synthetic and unmodified RNA control (scrambled). The latter was introduced to differentiate between proteins that were associating specifically with 5’ tsRNA-Glu^CUC/UUC^ and those that were *bona fide* RNA binders without specificity to 5’ tsRNA-Glu^CUC/UUC^. Label-free MS yielded a total of 1107 quantified proteins with at least one razor and unique peptide and a minimum of three LFQ values out of 12 experiments (supplementary Table 3). Two replicate experiments probing CPEs collected during steady-state conditions yielded 171 proteins with a positive fold change (log_2_ ratio ≧0) in both, and at least a 2-fold change (log_2_ ratio ≧1) in one of the replicates when normalized to noRNA control, and 168 proteins when normalized to the scrambled RNA control experiments (Figures 4C and D, supplementary Figures 4A and B, supplementary Table 3). Furthermore, LFQ analysis of peptides obtained from probing CPEs collected after iAs-exposure yielded 104 proteins when normalized to noRNA control, and 56 proteins when normalized to the scrambled RNA control experiments again with a fold change ≧1 in both (log_2_ ratio ≧0), and at least a 2-fold change (log_2_ ratio ≧1) in one of the replicates (Figures 4E and F, supplementary Figures 4A and B, supplementary Table 3). These data revealed specific protein associations with 5’ tsRNA-Glu^CUC/UUC^ but not with a generic RNA sequence and suggest that stress conditions might lead to changes in tsRNA-associated proteins.

**Figure 4.**
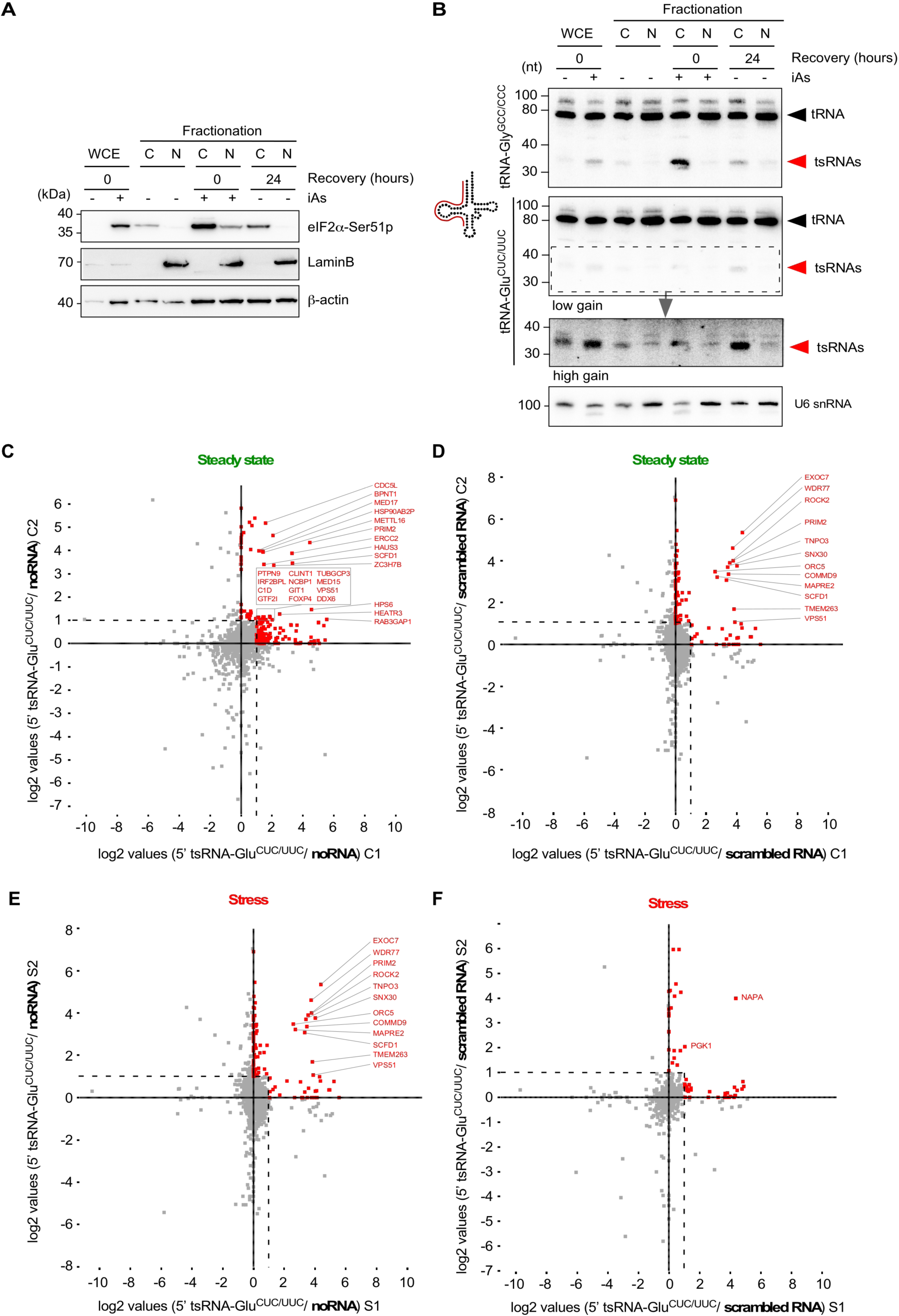
RNA affinity capture of proteins using purified tsRNAs. (A) Western blotting on fractionated HEK293T cell protein extracts obtained from control cells and cells that were exposed to 0.5 mM iAs for one hour followed by no or a recovery period of 24 hours before cell harvesting. Membranes were probed with antibodies against phosphorylated eIF2α and β-Actin as a loading control. (B) Northern blotting of total RNA (3 µg) extracted from fractionated HEK293T cell protein extract as described in (A) using 5’ probes against tRNA-Gly^GCC^ and tRNA-Glu^CUC^ (annealing of probe in full-length tRNA according to cartoon). Lower image for tRNA-Glu^CUC^ blot (high gain) represents a digitally enhanced region of the upper image (low gain) for better visualization of tsRNAs. Black arrowheads: full-length tRNAs; red arrowheads: tsRNAs. (C-F) Label-free quantitative comparison of protein enrichment in RNA affinity capture experiments using purified tsRNAs. (C) Scatter plot of protein ratios of replicate control HEK293T CPEs (steady state C1, C2) with 5’ tsRNA-Glu^CUC/UUC^ (purified from HEK293-ecANG cells) versus no RNA controls. (D) Scatter plot of protein ratios of replicate control HEK293 CPEs (steady state C1, C2) with 5’ tsRNA-Glu^CUC/UUC^ (purified from HEK293-ecANG cells) versus scrambled RNA controls. (E) Scatter plot of protein ratios of replicate iAs-exposed HEK293T CPEs (stress S1, S2) with 5’ tsRNA-Glu^CUC/UUC^ (purified from HEK293T-ecANG cells) versus no RNA controls. (F) Scatter plot of protein ratios of replicate iAs-exposed HEK293T CPEs (stress S1, S2) with 5’ tsRNA-Glu^CUC/UUC^ (purified from HEK293T-ecANG cells) versus scrambled RNA controls. A positive fold change (log_2_ ratio ≧ 1) when normalized to control (no or scrambled RNA) is indicated in red.

### Semi-quantification of tsRNA copy numbers per cell using purified tsRNAs

RNA modification-related biases in efficient reverse-transcription of tRNA-derived sequences, including various mapping issues, largely disqualifying sequencing-based methods from quantifying tRNA and tsRNA abundance^8–11^. Hence, the actual copy number of individual tRNAs or specific tsRNAs in any cell type under specific growth or stress conditions remains largely unknown. Microscale thermophoresis has been recently used to measure copy numbers of endogenous tRNAs^54^. As for tsRNAs, copy number estimates for particular tsRNA species have been attempted based on text book calculations (for instance, in supplementary data in^20^), but these estimates remain experimentally unproven. In an attempt to approximate the copy number of specific tsRNAs in iAs-exposed HEK293T cells, 5’ tsRNA-Gly^GCC/CCC^ and 5’ tsRNA-Glu^CUC/UUC^ levels were determined semi-quantitatively using northern blotting. To this end, total RNA extracted from a defined number of stressed cells was probed along with a dilution series of purified endogenously modified tsRNAs. Radiographic signals from serial tsRNA dilutions were blotted as standard curve and quantified to calculate the relative mass of individual stress-induced tsRNAs species per cell. When measuring the average yield of total RNA after Trizol extraction, the calculation indicated that this extraction method yielded about 4.75 picograms RNA from one million HEK293T cells (according to NanoDrop quantification). Blotting 4.5 μg of total RNA from HEK293T cells before and after iAs exposure against a dilution of purified 5’ tsRNAs resulted in signals that could be semi-quantitatively and comparatively quantified. Ensuing calculations indicated that the copy number of tsRNAs in a single HEK293T cell after iAs-induced stress was about 14.000 for 5’ tsRNA-Gly^GCC/CCC^ (Figures 5A and B) and about 12.000 for 5’ tsRNA-Glu^CUC/UUC^ Figures 5C and D).

**Figure 5.**
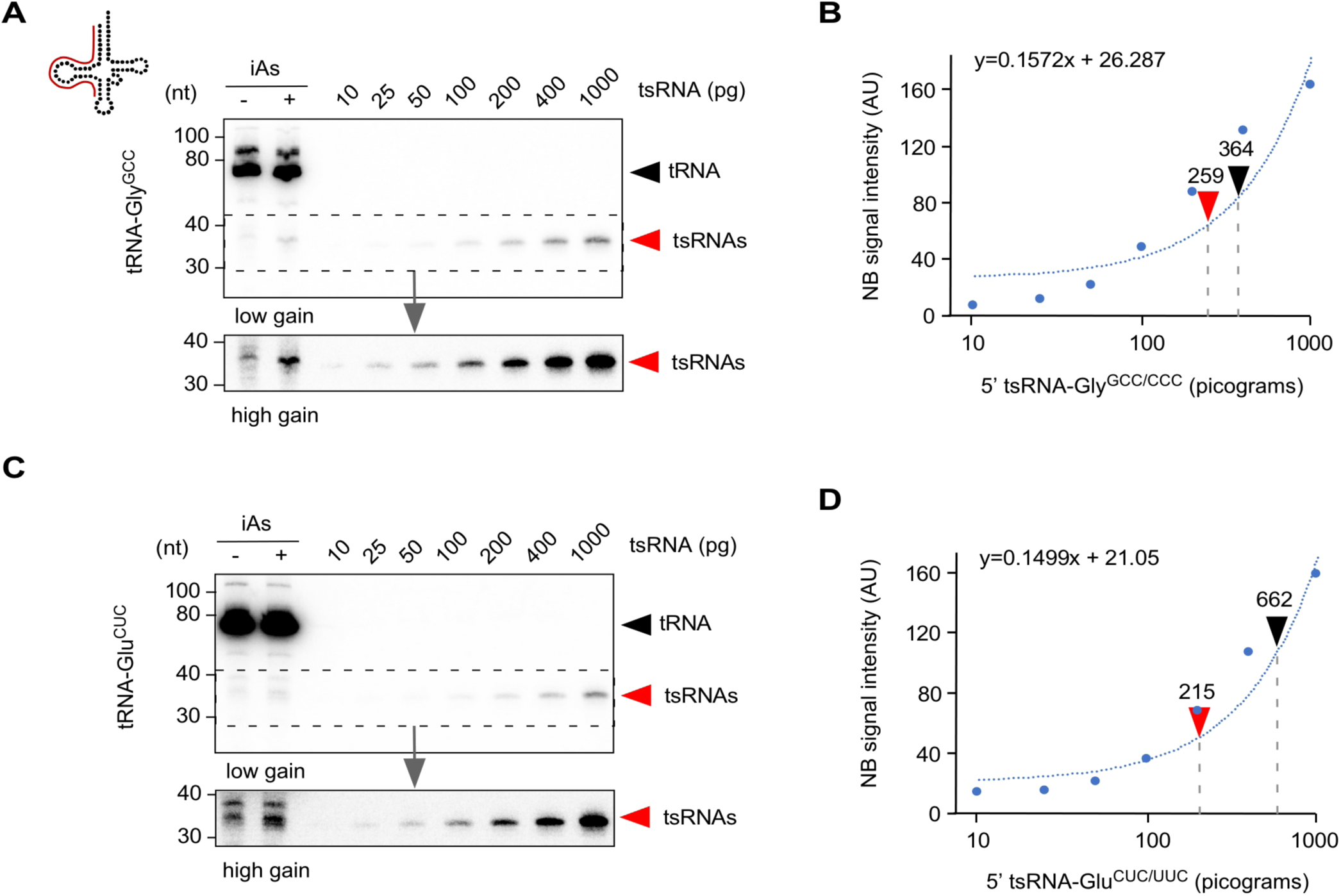
Semi-quantification of tsRNA numbers using purified tsRNAs. (A) Northern blotting of total RNA (4.5 µg) extracted from HEK293T cells under steady state conditions and after exposure to iAs along with a dilution series of purified 5’ tsRNA-Gly^GCC/CCC^ obtained from HEK293T-ecANG cells (72 hours constant Dox induction) using a 5’ probe against tRNA-Gly^GCC^ (annealing of probe in full-length tRNA according to cartoon). Lower image (high gain) represents a digitally enhanced region of the upper image (low gain) for better visualization of tsRNAs. Black arrowhead: full-length tRNAs; red arrowheads: tsRNAs. (B) Plotting of radiographic intensity values (arbitrary units, AU) derived from northern blotting of 5’ tsRNA-Gly^GCC/CCC^ as described in (A). tsRNA mass was logarithmically plotted on the x-axis (pictograms, pg). Trend line was used to derive mass values for radiographic intensities measured at the migration level of tsRNAs in control and iAs-exposed HEK293T cells (see A, high gain image). Black arrowhead: measured tsRNA mass without subtraction of control value; red arrowhead: measured tsRNA mass after subtraction of control value. (C) Northern blotting of total RNA (4.5 µg) extracted from HEK293T cells under steady state conditions and after exposure to iAs as described in (A) but probed with a 5’ probe against tRNA-Glu^CUC^ Lower image (high gain) represents a digitally enhanced region of the upper image (low gain) for better visualization of tsRNAs. Black arrowhead: full-length tRNAs; red arrowheads: tsRNAs. (D) Plotting of radiographic intensity values (arbitrary units, AU) for 5’ tsRNA-Glu^CUC/UUC^ as described in (A). A trend line was used to derive mass values for radiographic intensities measured at the migration level of tsRNAs in control and iAs-exposed HEK293T cells (see C, high gain image). Black arrowhead: measured tsRNA mass without subtraction of control value; red arrowhead: measured tsRNA mass after subtraction of control value.

## Discussion

tRNA-derived small RNAs (tsRNAs) can be extracted from many small RNA sequencing data sets. The biological significance of these small RNAs has been ignored until recently because their varying abundance and heterogeneity made them likely remnants of tRNA maturation or tRNA degradation intermediates. The potential for biological impact of specific tsRNAs has only been recognized after discovering that stress-induced tRNA fragmentation resulting in tRNA halves is a conserved part of the eukaryotic stress response with particular effects on protein translation and cellular survival^13, 19, 20, 55^. Ever since, an increasing number of reports has been assigning particular functions to specific tsRNAs suggesting a rather defined molecular impact on all kinds of cellular processes, which are not necessarily limited to the stress response. However, actual data on the mechanistic details as to how tsRNAs impact specific cellular processes remain scarce. For instance, it is unclear whether the few percent of isoacceptor-specific tsRNAs act as single entities or rather in bulk with tsRNAs derived from other tRNAs. Furthermore, many of the current reviews re-iterate mere correlations thereby connecting tsRNA detection and abundance (mostly after RNA sequencing) with a wide range of cellular pathways, often without mentioning that the mechanistic underpinnings of tsRNA function have often not been addressed yet (critically discussed in^1^).

To better understand the mechanistic details of tsRNA function, improved methodology needs to be developed, which would allow measuring actual tsRNA copy numbers, localizing specific tsRNAs *in situ*, mapping tsRNA modification patterns and calculating their stoichiometry, as well as determining the molecular interactions of specific tsRNAs. The experimental basis for addressing most of these questions should be the availability of pure tsRNA sequences, preferably from cellular sources where tsRNAs are being produced. This is important because tRNAs are the most highly modified tRNAs in any cell type, making it likely but largely untested that the functionality of specific tsRNAs depends on their modification status. Biochemical attempts for enriching small RNAs including tsRNA have been published^56^. However, reports that aimed at addressing function or molecular interactions of specific tsRNAs have almost exclusively used synthetic RNA sequences (an exception being^57^) thereby largely ignoring the possibility that tsRNA structure and function could be determined by specific RNA modifications.

Theoretically, chemical RNA synthesis allows introducing modified nucleotides at specific positions given that these positions are known once such modification patterns have been determined. However practically, many modified nucleotides remain commercially unavailable, which necessitates often complicated chemical synthesis by expert laboratories. Furthermore, commercially available (unmodified) RNAs appear to contain trace amounts of modified nucleotides^58^, which might affect experimental outcomes when testing synthetic or synthetically modified tsRNAs for structure-function relationships.

A viable alternative is the purification of tsRNAs from endogenous sources and under conditions that promote tRNA fragmentation. Importantly, specific tRNAs and their modification patterns have been systematically determined using hybridization-based affinity capture of target RNAs from complex samples followed by LC-MS/MS^59^.

Here, we used similar methodology to purify specific tsRNAs from human cells after exposure to tRNA fragmentation-inducing stress conditions or when ectopically expressing the mammalian anticodon nuclease Angiogenin. Our results suggest that these *in vivo* and *in vitro* manipulations could produce any tsRNA species in scalable fashion, which can be used to address a variety of open questions relating to tsRNA sequences. For instance, purified tsRNAs containing specific modification patterns could be used for *in vitro* protein capture experiments, which should be compared to experiments with synthetic tsRNA sequences thereby addressing the modification-specific association of proteins with RNAs. Furthermore, purified tsRNAs containing specific modification patterns could be used for amplification-independent tsRNA quantification, which, in combination with using synthetic tsRNA sequences, might allow determining the effects of RNA modifications on hybridization-based read-outs. The calculated values fit well to previous calculations as to how many individual tsRNAs might exist in a single cell (supplemental information in^20^) although all quantifications are dependent on the initial measurements of RNA concentrations, the outcome of which can differ by magnitudes depending on the quantification method used^60, 61^. In addition, the recently reported possibility of intra-and intermolecular cross-hybridization between 5’ tsRNA-Gly^GCC^ and 5’ tsRNA-Glu^CUC^ in extracellular space^62^ could be tested for modification-dependence of such hybrid formation using purified tsRNAs and after determining the RNA modification patterns of secreted tsRNAs. Furthermore, reported tsRNA activities as *bona fide* small RNA entities with posttranscriptional gene regulatory functions akin to siRNAs or miRNAs^30, 35^ need to be tested with modified tsRNAs since various RNA modifications affect the base-pairing capabilities of small RNAs and therefore might modulate the activities that were originally reported for synthetically introduced tsRNAs. Lastly, the recently reported effect of RNA modifications on the efficiency of sperm-borne small RNAs (including tsRNAs) for intergenerational transmission of paternally acquired metabolic disorders^63, 64^ raises the question as to which RNAs exactly contribute to these processes. The present consensus points towards a role for sperm-carried tsRNAs but the identity, exact modification status of individual tsRNA species and their abundance has not been addressed in molecular detail. Taken together, the presented workflow provides the basis for the systematic purification of specific and modified tsRNAs which could be used for probing various unresolved aspects in tsRNA biology.

## Supporting information

STable 1

STable 2

STable 3

## Additional information

## Acknowledgements

This work was supported by the Austrian Science Foundation (FWF-P29094). AD is a fellow of the ÖAW DOC graduate program. Stefanie Kellner is funded by Deutsche Forschungsgemeinschaft (DFG), grant number: KE1943/3-1. Proteomics analyses were performed in the framework of the Vienna Life Science Instruments (VLSI) initiative at the Max Perutz Laboratories Mass Spectrometry Facility using the VBCF instrument pool.

## Contributions

MS and AD conceived experiments and wrote the manuscript. AD established and characterized HEK293T-ecANG cells, established the methods for chromatography and affinity capture, performed northern and western blotting on ecANG experiments. AD and VO performed tRNA and tsRNA purifications. VO performed northern blotting for iAs-induced tRNA fragmentation. S. Kellner and S. Kaiser processed tRNA and tsRNA samples for LC-MS/MS, performed mass spectrometry and analyzed the data. CT and EJ performed ecANG purification and ribozyme-mediated *in vitro* transcription of synthetic tsRNAs. CT performed ecANG on beads fragmentation of tRNAs, designed DNAzymes, performed *in vitro* cleavage assays and measured cell viability and tsRNA abundance. DA and MH performed analysis of protein mass spectrometry data and provided visualization of data.

## Disclosure statement

No potential conflict of interest was reported by the author(s).

## Accession numbers

Small RNA sequencing has been deposited at NCBI GEO database under accession number GSE139124.

## Supplementary Figure legends

**Supplementary Figure 1.**
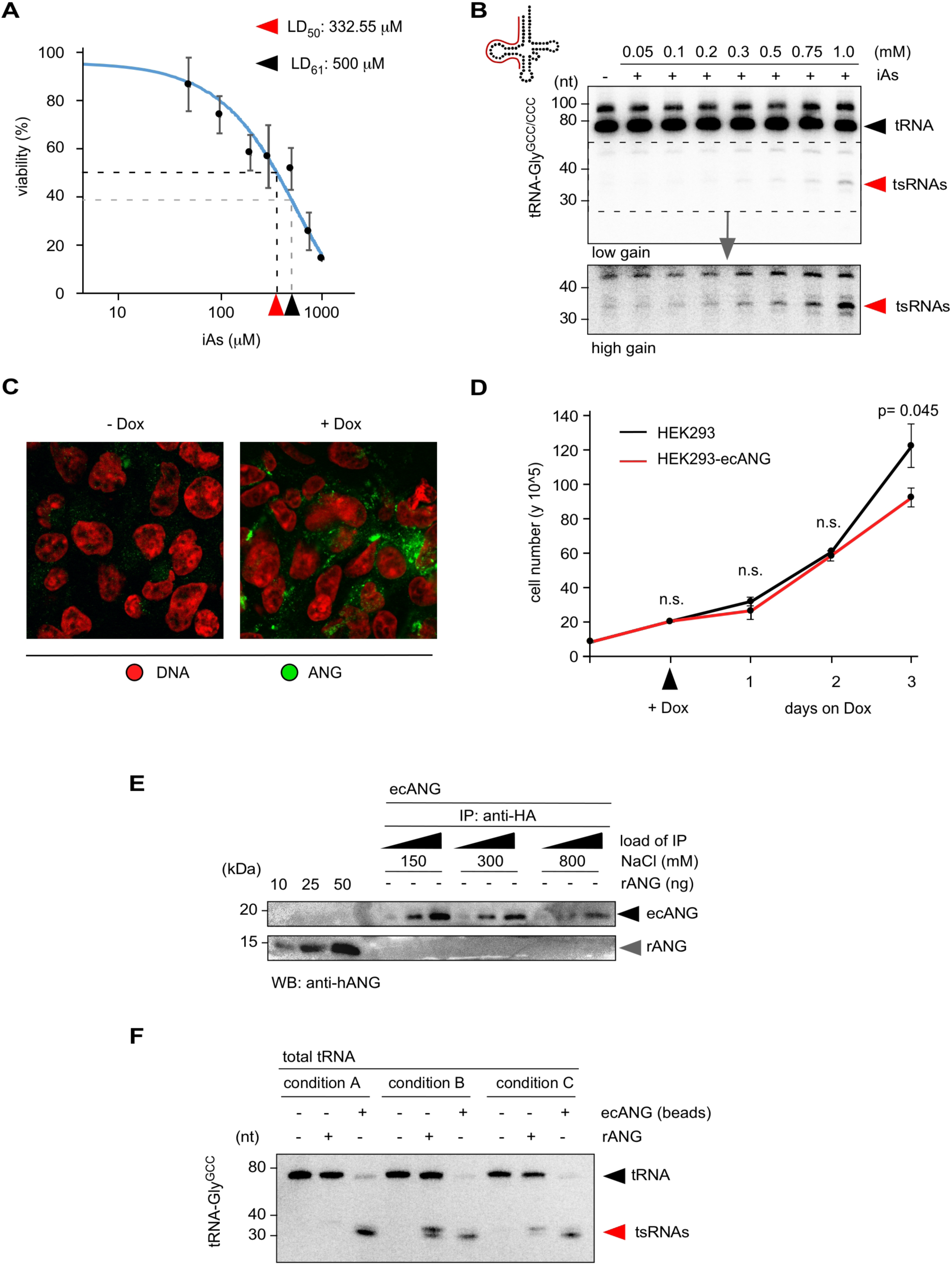
Comparison of cell viability during tsRNA production. (A) Lethal dose (LD_50_) determination of HEK293T cells after exposing to different iAs concentrations. Triplicate cell viability measurements 24 hours after iAs exposure and wash-out indicate an LD_50_ value of 333 µM (red arrowhead), a concentration of iAs that is still below the commonly used iAs concentration (500 µM, black arrowhead), which would correspond to a LD_61_ value. (B) Northern blotting of total RNA (3 µg) from HEK293T cells exposed to different iAs concentrations (LD_20_ to LD_60_) using 5’ probes against tRNA-Gly^GCC^ and tRNA-Glu^CUC^ (annealing of probe in full-length tRNA according to cartoon). Black arrowhead: full-length tRNAs; red arrowhead: tsRNAs. (C) Indirect immuno-fluorescence images of HEK293T-ecANG cells before and after ecANG expression using Dox using antibodies against human ANG. Red: DNA; green: ANG. (D) Triplicate cell viability measurements of HEK203T-ecANG cells over the course of Dox-mediated ecANG expression. Statistical analysis was performed using a Student’s t-test (equal variance). (E) Western blotting on a dilution series of recombinant ANG and immuno-precipitated ANG from HEK293T-ecANG-derived cell culture medium (1 mL collected 48 hours after Dox-induction) using antibodies against human ANG (hANG). Immuno-precipitated ecANG on beads was washed with buffers of different stringency (NaCl concentration) and different volumes of the immuno-precipitated eluate were analysed to determine the mass of ecANG that can be precipitated from a given volume of cell culture medium. (F) Northern blotting on recombinant ANG-mediated or ecANG-mediated *in vitro* tRNA fragmentation after purification of total tRNAs under native conditions using acidic phenol and native gel extraction followed by no melting and re-annealing of tRNA fraction (condition A) or melting and reannealing of tRNA fraction (condition B) or purification of total tRNA using Trizol followed by denaturing gel elution, melting and reannealing of tRNA fraction (condition C). The blot was hybridized with a 5’ probe against tRNA-Gly^GCC^. Black arrowhead: full-length tRNAs; red arrowhead: tsRNAs.

**Supplementary Figure 2.**
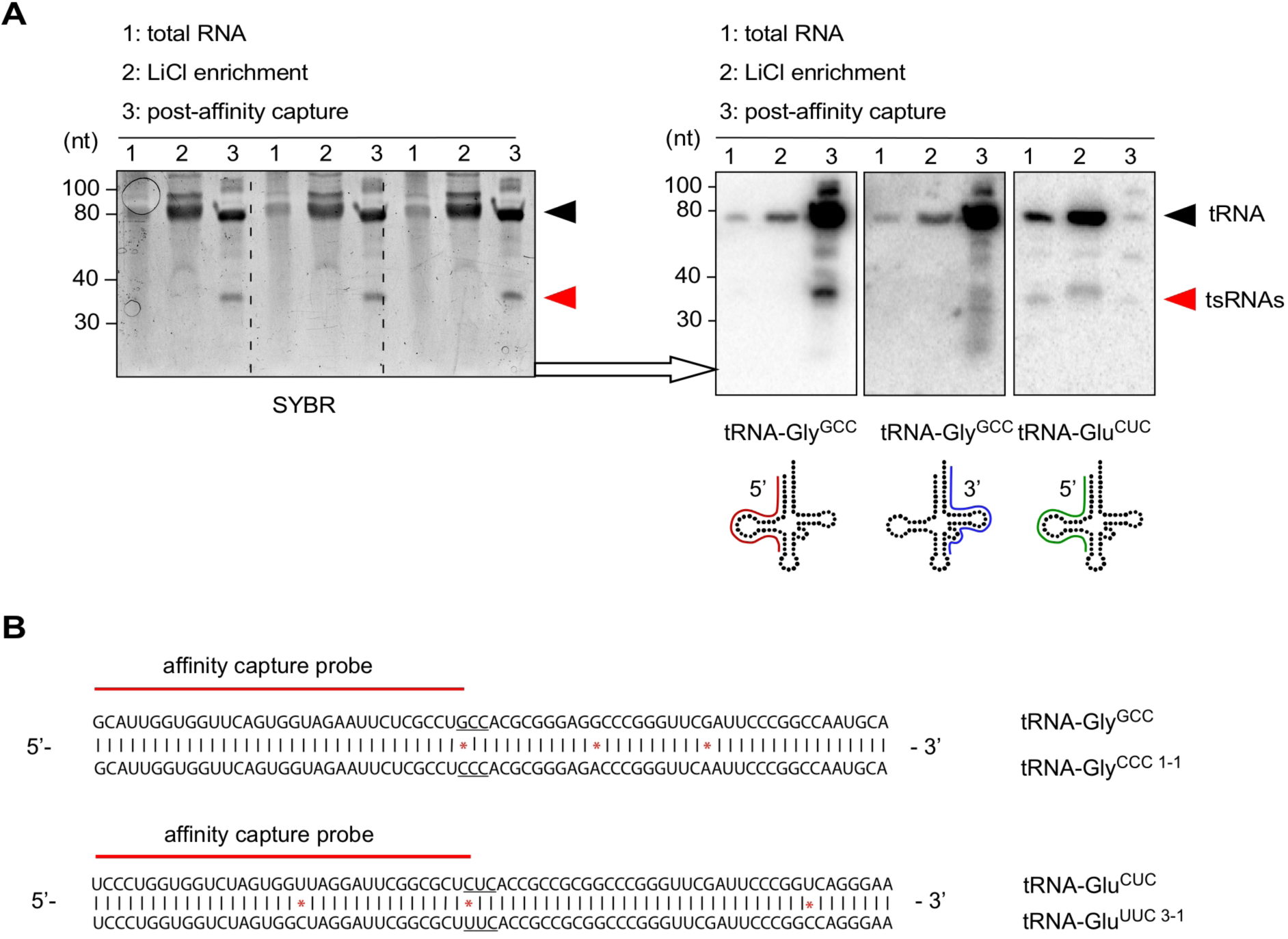
(A) SYBR staining of urea-PAGE after small RNA enrichment using LiCl precipitation (LiCl enrichment) and affinity capture of tRNA-Gly^GCC^-derived sequences from the LMW RNA pool (in triplicate). Northern blotting of total RNA, LiCl-enriched LMW RNA and affinity captured tRNA- and 5’ tsRNA-Gly^GCC^ using probes against the 5’ portion of tRNA-Gly^GCC^, the 3’ portion of tRNA-Gly^GCC^ and the 5’ portion of tRNA-Glu^CUC^. Black arrowheads: full-length tRNAs; red arrowheads: tsRNAs. (B) Sequences of full-length tRNA-Gly^GCC^, tRNA-Gly^CCC-1.1^, tRNA-Glu^CUC^ and tRNA-Glu^UUC-3.1^(according to http://gtrnadb.ucsc.edu/), which could be captured by the used NHS-coupled complementary oligonucleotides (depicted in red).

**Supplementary Figure 3.**
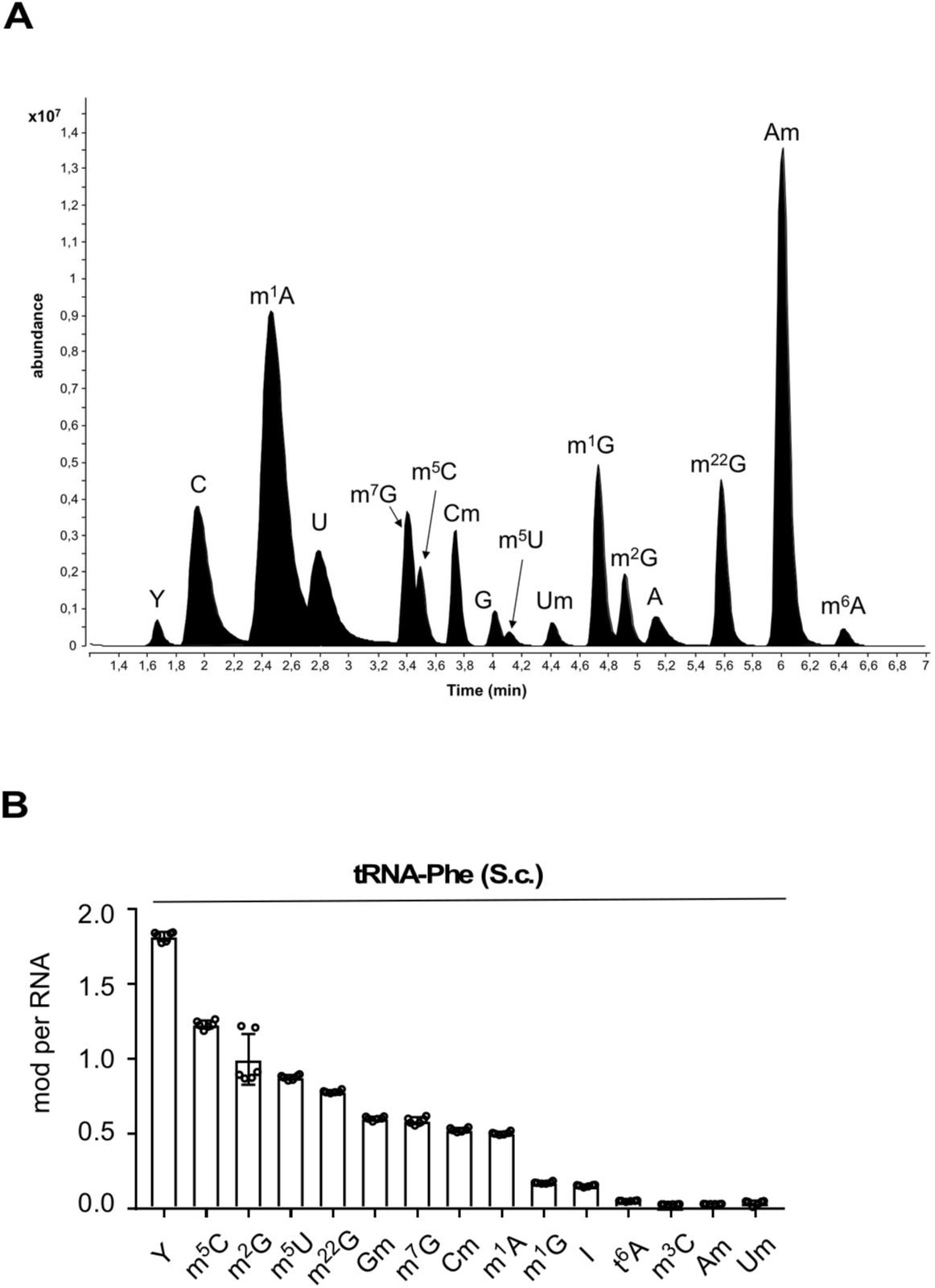
(A) LC-MS/MS chromatogram of an exemplary calibration measurement preceding the analysis of tRNA-derived sequences. (B) Results from sextuplicate LC-MS/MS analysis of tRNA-Phe purified from yeast (*Saccharomyces cerevisiae*. S.c.). From six replicate measurements, plotted as single data points with mean and standard deviation. Abbreviations: Y (pseudouridine), C (cytidine), m^1^A (1-methyladenosine), U (uridine), m^7^G (7-methylguanosine), m^5^C (5-methylcytidine), m^3^C (3-methylcytidine), Cm (2’-O-methylcytidine), G (guanosine), Gm (2’-O-methylguanosine), m^5^U (5-methyluridine), Um (2’-O-methyluridine), m^1^G (1-methylguanosine), m^2^G (2-methylguanosine), A (adenosine), I (inosine), m^22^G (2,2-dimethylguanosine), Am (2’-O-methyladenosine), m^6^A (6-methyladenosine), t^6^A (N6-threonylcarbamoyladenosine).

**Supplementary Figure 4.**
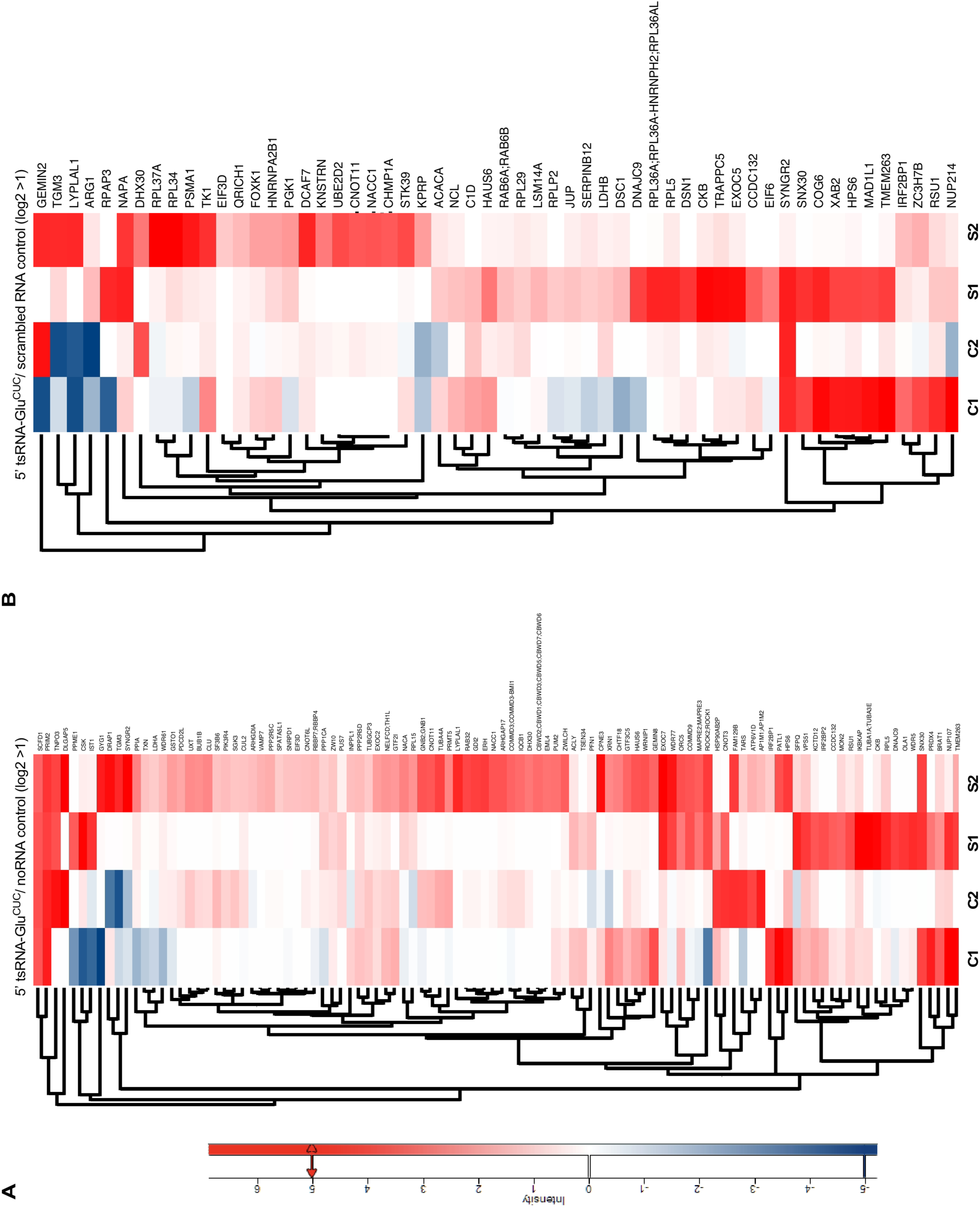
(A) Heatmaps depicting protein associations with 5’ tsRNA-Glu^CUC/UUC^ (purified from HEK293T-ecANG cells) for two replicate experiments on control HEK293T CPEs (C1, C2) and CPEs extracted from iAs-exposed HEK293T cells (S1, S2) when normalized to noRNA controls. For stress-control comparison, protein hits with a positive log-fold change in both replicates and a log-fold change ≧ 1 in at least one of the two replicates under stress conditions were selected and represented in maps. For actual log_2_ ratios, see supplementary Table 3. (B) Heatmaps depicting protein associations with 5’ tsRNA-Glu^CUC/UUC^ (purified from HEK293T-ecANG cells) for the same experiments as in (A) but normalized to scrambled RNA controls. For stress-control comparison, protein hits with a positive log-fold change in both replicates and a log-fold change 1 in at least one of the two replicates under stress ≧ conditions were selected and represented in maps. For actual log_2_ ratios, see supplementary Table 3.

## Supplementary Table legends

**Supplementary Table 1**

Dynamic multiple reaction monitoring (MRM) parameters for the detection and quantification of modified nucleosides.

Abbreviations: MS1 Res, MS1 resolution; MS2 Res, MS2 resolution; Ret Time, retention time (with a one minute retention time window); SILIS, stable isotope-labelled internal standard (as described in^65^).

**Supplementary Table 2**

Results from biological triplicate experiments measuring tRNA- and 5’ tsRNA-Gly^GCC/CCC^ and tRNA- and 5’ tsRNA-Glu^CUC/UUC^ either after iAs exposure or ecANG expression. Absolute abundance of modified nucleosides per purified RNA (tRNA and 5’ tsRNAs derived from tRNA-Gly^GCC/CCC^ and tRNA-Glu^CUC/UUC^) was determined by isotope dilution mass spectrometry (as described in^65^).

**Supplementary Table 3**

List of proteins identified by mass spectrometry analysis of RNA affinity purified CPEs filtered for a minimum of three LFQ values over 12 LC-MS/MS experiments. Proteins enriched by endogenous RNA (endo) versus no RNA and scrambled RNA (controls) from stressed and steady-state conditions are listed separately based on the LFQ intensities ratios showing a positive log-fold change in both replicates and a log-fold change D 1 in at least one of the two replicates.

## Materials and methods

### Cell culture

HEK293T cells were cultured in standard Dulbecco’s Modified Eagle’s Medium (DMEM, Sigma Aldrich) supplemented with penicillin (100 U/mL), streptomycin (100 μg/mL), 2 mM L-μ glutamine and 10 % fetal bovine serum in a humidified incubator at 37°C and 5 % CO_2_.

### Determining LD_50_ for iAs exposure

HEK293T cells were cultured to 70 % confluency, treated for one hour with different concentrations of inorganic sodium arsenite (iAs) followed by washout of iAs and re-plating in fresh medium. After 24 hours, cells were harvested by trypsinization, stained with Trypan-Blue and counted using a Neubauer-chamber.

### Stress experiments

For oxidative stress, iAs was added at stated final concentrations to HEK293T cells at 70 % confluency for one hour, followed by washout and re-plating until harvest.

### Ectopic human ANG expression system

A HEK293T cell line harboring an inducible expression cassette containing human Angiogenin-HA-FLAG (ecANG) was established after co-transfection of the ecANG destination vector and the pOG44 recombinase expression plasmid followed by clonal selection of cells transfected cells. This cell line was cultured under selection using 500 μg/mL Hygromycin B (stock: 100 mg/mL) and 75 μg/mL Blasticidin (stock: 10 mg/mL). At 70-80 % confluency, ecANG expression was induced by adding doxycycline (Dox) to 1μg/mL. Cells were cultured in the continuous presence of Dox for up to 72 hours.

### Determining cell viability during Dox-mediate ecANG expression

HEK293T-ecANG cells were cultured to 50 % confluency, doxycycline was added to 1μg/mL and living cells were counted every day for three days using Trypan-Blue.

### Whole cell protein extraction

For harvesting by scraping, cell culture media was removed from the cells and ice-cold 1x PBS was added. Cell pellets were homogenized in PD Buffer (20 mM Tris pH 7.4, 150 mM NaCl, 10 mM MgCl_2_, 10 % (v/v) Glycerol, 0.2 % (v/v) NP-40, 1x protease inhibitor cocktail, Roche) using a 26G needle (Sterican) followed by two centrifugations for 10 minutes at full speed at 4°C.

### Sub-cellular fractionation

HEK293T cells were harvested into ice-cold 1x PBS. Cells were gently homogenized in CE-B (150 mM NaCl, 50 mM HEPES pH 7.4, 25 μg/ml digitonin, 1x protease inhibitor cocktail) for 10 minutes at 4°C under constant rotation followed by a centrifugation for 10 minutes at 2.000 x g and 4°C. The 2.000 x g supernatant was centrifuged an additional 10 minutes at full speed and 4°C, after which sediment was discarded and the remaining supernatant was considered as the soluble cytoplasmic fraction. The 2.000 x g pellet was washed in-ice cold 1x PBS followed by homogenization in ice-cold MOE-B (150 mM NaCl, 50 mM HEPES pH 7.4, 1 % (v/v) NP-40, 1x protease inhibitor cocktail). Homogenates were incubated on ice for 30 minutes followed by centrifugation at 7.000 x g for 10 minutes to pellet nuclei. Nuclei were washed in ice-cold 1x PBS and homogenized in NE-B (20 mM Tris-HCl pH 8.0, 420 mM NaCl, 1.5 mM MgCl_2_, 0.2 mM EDTA, 1 mM PMSF, 25 % (v/v) Glycerol, 1x protease inhibitor cocktail). Nuclei were extracted by rotation at 4°C overnight and the soluble nuclear protein fraction was collected by centrifugation for ten minutes at 7.000 x g at 4°C.

### Western blotting

For whole protein lysates of HEK293T cells, a cell pellet was homogenized in RIPA buffer (150 mM NaCl, 1 % (v/v) NP-40, 0.1 % (v/v) SDS, 50 mM Tris, pH 8.0, 0.5 % (v/v) sodium deoxycholate, 1x protease inhibitor complex, Roche). Samples were incubated on ice for 10 minutes, followed by two centrifugations, 15 minutes each at 16.000 x g. 50 μ of total protein extracts were solubilized in SDS-sample buffer by boiling at 95°C for 5 minutes and analyzed by SDS-PAGE. Western blotting was performed with antibodies against human ANG (goat, R&D systems; 1:300), β-actin (rabbit, SIGMA; 1:1.000), eIF2α-P (rabbit, Abcam 32157; 1:500), HA (mouse 16B12, Covenance; 1:1.000), laminB (goat, Santa Cruz; 1:5.000).

### RNA extraction

Collected cell pellets were re-suspended in 1x PBS and RNA was extracted using self-made Trizol solution (38 % (v/v) phenol, 800 mM guanidine thiocyanate, 400 mM ammonium thiocyanate, 100 mM NaOAc, 5 % (v/v) glycerol, 0.5 % (w/v) N-lauroylsarcosine). Samples were incubated at room temperature for five minutes, followed by chloroform extraction and isopropanol precipitation. RNA pellets were washed in 75 % ethanol and re-suspended in RNase-free water.

### Northern blotting

RNA was separated on 12 % Urea-PAGE in 0.5x TBE and transferred to Nylon membranes (Roche, GE Healthcare) using semi-dry blotting in 0.5x TBE for 30 minutes at 5 V=constant. Transferred RNA was immobilized by UV cross-linking (StrataLinker), followed by hybridization with ^32^P-end labeled oligonucleotides in blocking solution (5x SSC, 20 mM Na_2_HPO_4_ pH 7.4, 1 % SDS, 1x Denhardt’s reagent) at 39°C overnight. After washing with 3x SSC, 5 % (v/v) SDS for 15 minutes and 3x SSC, 5 % (v/v) SDS for 10 minutes, membranes were exposed at room temperature to a magnetic screen and imaged using an Amersham Typhoon Biomolecular Imager (GE-Healthcare).

### Differential precipitation of small RNAs using Lithium chloride

Harvested cell cultures were processed to separate high-molecular weight (HMW) from low-molecular weight (LMW) RNAs as described in^66^ but without pre-treatment steps. Instead total RNA was extracted using Trizol followed by LiCl precipitation of HMW RNAs overnight. The resulting soluble RNA content was precipitated with 1/10 volume of 3 M sodium acetate (pH 5.2) and 2.5 volume of pre-cooled absolute ethanol at −20°C overnight.

### tRNA and 5’ tsRNA isolation from HEK293 cells

Total RNA was isolated from five 15 cm ⊘ cell culture plates exposed to 0.5 mM iAs for one hour or from ecANG-expressing HEK293T cells after three days of Dox induction. In parallel, tRNAs from iAs-exposed cells were extracted. tsRNAs from iAs-exposed cells and ecANG-expressing cells were purified as follows. Total RNA was isolated from HEK293T cells using Trizol. Total RNAs extracted from five 15 cm ⊘ cell culture plates were re-suspended in 10 mL of IEX-buffer (20 mM Tris, 10 mM KCl, 1.5 mM MgCl_2_). For ion exchange chromatography, a HiTrap Q FF anion exchange chromatography column (1 mL, GE Healthcare) was used on an ÄKTA-FPLC (GE Healthcare) at 4°C. For elution, NaCl was used as eluent. Eluted fractions between 400-600 mM NaCl were collected and immediately precipitated using isopropanol at −20°C. RNA was re-suspended, supplemented with 10 mM MgCl_2_ and renatured by incubation at 75°C for three minutes. Afterwards the RNA was immediately placed on ice and added to 20 ml NHS-BB (30 mM HEPES KOH, 1.2 M NaCl, 10 mM MgCl_2_). For RNA affinity capture, 80 µg of a 5’ amino-modified DNA oligonucleotide complementary to the target tRNA (IDT) was covalently coupled to a HiTrapTM NHS-activated HP column (1 mL, GE Healthcare). For isolation of full-length tRNAs, binding to the NHS-column was performed in a touch-down style. The small RNA pool was circulated for 90 minutes at 65°C and the temperature was gradually decreased over a three-hour period until reaching 40°C and thereafter washed using NHS-WBA (2.5 mM HEPES KOH, 0.1 M NaCl, 10 mM MgCl_2_). For isolation of 5’ tsRNAs, the small RNA fraction in NHS-BB was circulated through the column for four hours at 60°C and was afterwards washed using NHS-WBA. Bound RNAs were eluted by submerging the column in a water bath at 75°C in NHS elution buffer (0.5 mM HEPES-KOH, 1mM EDTA) followed by immediate precipitation in isopropanol at −20°C. To maintain column for reuse, it was washed with three volumes of NHS-SB (0.05 M Na_2_HPO_4_, 0.1 % (w/v) NaN_3_, pH 7.0) and stored at 4°C. Precipitated RNA was re-suspended in water. Purified tRNA-Gly^GCC^ was further gel-purified (8 % urea-PAGE in 0.5 x TBE) using RNA gel extraction buffer (0.3 M NaOAc, pH 5.2, 0.1 % (v/v) SDS, 1 mM EDTA) and immediately precipitated in isopropanol at −20°C. Precipitated RNA was re-suspended in water.

### Isolation of ecANG by immuno-precipitation

ecANG was isolated from cell culture supernatants of Dox-induced HEK293T-ecANG cells. 72 hours post-induction, supernatants were harvested and cleared of cells and debris by centrifugation. Dox-induced HEK293T cell supernatant (one mL) was incubated with 50 µL anti-FLAG-M2 magnetic beads (50 % suspension, SIGMA) for one hour at room temperature, followed by three washing steps in PBS or buffers containing increasing concentration of NaCl. The amount of isolated ecANG was measured against defined masses of recombinant ANG using western blotting. Immuno-precipitated ecANG on beads was directly used for *in vitro* cleavage of tRNAs.

### Recombinant ANG-and ecANG-mediated tsRNA production

For cleavage reactions using recombinant ANG (R&D Systems), 500 ng of total RNA or purified tRNA-Gly^GCC^ were melted for 5 minutes at 70 °C in H_2_O, put on ice for 5 minutes followed by addition of cleavage buffer (30 mM HEPES pH 6.8, 30 mM NaCl, 10 mM MgCl_2_). The reaction was incubated for 10 minutes at room temperature before adding either 100 ng of recombinant ANG to a final reaction volume of 15 µL or the reaction was added to ecANG on beads (from one milliliter cell culture supernatant) followed by incubation at 37°C for two hours. The reaction was separated from magnetic beads and used for further analysis.

### DNAzyme-mediated tsRNA production

500 ng of purified tRNA-Gly^GCC/CCC^ was incubated in a volume of 15 µl with a 10-fold excess of DNAzyme in 50 mM Tris-HCl (pH 7.5), 150 mM KCl, 10 mM MgCl_2_ (as described in ^48^). DNAzyme reactions were performed with 30 iterations of: denaturing step at 85°C for 30 seconds followed by incubation at 37°C for three minutes.

### Separation of residual tRNAs from tsRNA isolates

To separate residual co-purified tRNAs from tsRNAs, precipitated RNAs were subjected to size exclusion chromatography. After re-suspending the affinity purified RNA in water in a total volume of 50 μL, the RNA was separated using an Advance Bio SEC 130A column (Agilent) in SEC buffer (20 mM Tris pH 7.4, 0.15 M NaCl) under a 0.75 mL/min flow rate. Fractions of 0.75 mL were collected and precipitated with isopropanol at −20°C.

### Removal of residual tRNAs from tsRNA isolates using RNase H and gel purification

To remove residual co-purified tRNAs from tsRNAs, precipitated RNAs were mixed with equimolar amounts of DNA oligonucleotides complementary to the 3’ half of the respective tRNA. The mix was heated to 80°C for three minutes, followed by reverse transcription using Superscript III (ThermoFisherScientific). In order to maintain tRNA-cDNA hetero-duplexes, final step of 85°C was omitted and the reaction was kept on ice. The reaction was supplemented with RNase H buffer and RNase H (10 U) and incubated at 37°C for 30 minutes for targeted degradation of tRNA. Exonuclease I (20 U, NEB) were added to digest cDNA and oligonucleotide sequences. Remaining tsRNAs were purified by acidic phenol-chloroform treatment and precipitation in isopropanol.

### Small RNA library preparation and cDNA sequencing

100 ng of purified tsRNAs were treated with 0.5 U/µL T4 polynucleotide kinase (NEB) and 166 µM ATP (NEB) for 40 minutes at 37°C. After clean-up using acidic phenol-chloroform extraction followed by isopropanol precipitation, small RNA library preparation was performed using the NEBNext Multiplex Small RNA Library Prep Set for Illumina (Set 2) (NEB). cDNA libraries were sequenced on a HiSeq2000 platform in a spike-in format in paired-end 50 (PE50) mode.

### Small RNA sequencing analysis

Small RNA reads were de-multiplexed, quality-controlled (fastqc) and trimmed (Trimmomatic) to remove adapters and barcodes, and to trim low quality bases. Reads were mapped to the human genome (release hg19) using Bowtie2 and mapping parameters: -N 0 -L 22. Output alignment files were used to intersect with the coordinated of the predicted tRNA annotation in ENSEMBL and counted as total number of reads mapping to an annotated tRNA gene.

### LC-MS/MS analysis of RNA modifications

About 400 ng of purified RNA was digested using a mixture of benzonase (2.5 U), bacterial alkaline phosphatase (10 U) and phosphodiesterase I (0.1 U) in a final reaction volume of 20 µL. The reaction mixture was supplemented with MgCl_2_ to a final concentration of 1 mM and Tris-HCl (pH 8.0) to a final concentration of 50 mM. Nucleobase deaminase inhibitor coformycin and tetrahydrouridine were added at a concentration of 10 μg/mL and 50 µg/ml, respectively, and butylated hydroxytoluene (an antioxidant) was added at a concentration of 0.5 mM (for further detail see^67^). The digestion was allowed to proceed for 2 hours at 37°C and was stopped by filtering through a 10 kDa MWCO filter (AcroPrepTM Advance, 350 µl, OmegaTM 10K MWCO, Pall, Dreieich, Germany) at 3000 x g for 30 minutes. After addition of 10 µl pure water for salt dilution purposes, 18 µL of filtrate was mixed with 2 µL of internal standard (produced as recently described^68^). 10 µL of each sample was injected for LC-MS/MS analysis (corresponding to around 150 ng tRNA digest). Calibration solutions for absolute quantification were prepared as recently described^68^. For quantification, an Agilent 1290 Infinity II equipped with a DAD combined with an Agilent Technologies G6470A Triple Quad system and electro-spray ionization (ESI-MS, Agilent Jetstream) was used. Operating parameters were as follows: positive ion mode, skimmer voltage 15 V, Cell Accelerator Voltage 5 V, N_2_ gas temperature 230°C and N_2_ gas flow 6 L/min, sheath gas (N_2_) temperature 400°C with a flow of 12 L/min, Capillary Voltage of 2500 V, Nozzle Voltage of 0 V and the Nebulizer at 40 psi. The instrument was operated in dynamic MRM mode and the individual mass spectrometric parameters for the nucleosides are given in supplementary Table 1. The mobile phases were: (A) as 5 mM NH OAc (≥ 99 %, HiPerSolv CHROMANORM^®^, VWR) aqueous buffer, brought to pH = 5.6 with glacial acetic acid (99 ≥ %, HiPerSolv CHROMANORM^®^, VWR) and (B) as pure acetonitrile (Roth, LC-MS grade, purity: 99.95 %). A Synergi Fusion-RP column (Phenomenex^®^, Torrance, California, USA; Synergi^®^ 2.5 µm Fusion-RP 100Å, 150 x 2.0 mm) at 35°C and a flow rate of 0.35 mL/min was used. The gradient began with 100 % A for one minute, increased to 10 % B by five minutes, and to 40 % B by 7 minutes. The column was flushed with 40 % B for one minute and returned to starting conditions to 100 % A by 8.5 minutes followed by re-equilibration at 100 % A for 2.5 additional minutes.

### In vitro transcription of tsRNA sequences using hammerhead ribozymes

Single-stranded DNA templates (IDT) encompassing a T7 promoter followed by hammerhead ribozyme and the specific 5’ tRNA isoacceptor or scrambled RNA sequences were PCR-amplified and afterwards *in vitro*-transcribed using 1.875 U/µL T7 Polymerase (NEB) in 1x T7-Buffer with 20 mM DTT, 18 mM spermidine, 3 µL murine RNase Inhibitors (40 U/µL NEB) and 125 µM NTP-mix (NEB). Transcribed and processed products were size selected on 8 % urea-PAGE and subsequent extraction using RNA gel extraction buffer (0.3 M NaOAc (pH 5.4), 0.1 % (v/v) SDS, 1 mM EDTA).

### Biotinylation of small RNAs

5’ tsRNA-Glu^CUC^ was de-phosphorylated using 0.5 U/µg Fast-AP Thermosensitive Alkaline Phosphatase (ThermoFisher Scientific) in 1x Fast-AP Buffer for 12 minutes at 37°C followed by five minutes at 75°C. RNA was isolated using acid Phenol/Chloroform/Isoamylalcohol (P/C/I pH 4.5, ROTH) and precipitated using isopropanol. De-phosphorylated tsRNA or synthetic RNA (scrambled) were 5’-thiolated by incubation with 0.5 U/µL polynucleotide   kinase (ThermoFisher Scientific) in the presence of 0.5 mM ATPγS (SIGMA) at 37°C overnight. EDTA was added to 1 mM final, PNK was inactivated by incubation at 75°C for five minutes and RNA was isolated using P/C/I, followed by precipitation in isopropanol. In each biotinylation reaction, one µg of small RNAs was mixed with 2 µg of HPDP-Biotin (ThermoFisher Scientific, 1 mg/ml in DMSO). The reaction was incubated for three hours at room temperature protected from light and mixed every 15 minutes. To monitor successful biotinylation, an increase in absorbance at 343 nm, which reports on the accumulation of a reaction by-product, pyridine-2-thione as a proxy for the biotinylation efficiency, was measured using NanoDrop (ThermoFisher Scientific). Biotinylated small RNAs were separated from HPDP-biotin and pyridine-2-thione using spin columns (BioRad) in ultra-pure water.

### RNA affinity capture

The RNA solution was supplemented to 10 mM MgCl_2_, denatured for three minutes at 75°C and cooled down to room temperature to re-nature the RNA. 450 µg CPE was pre-cleared by addition of 20 µl packed streptavidin-sepharose beads and incubation for one hour at 4°C under rotation. 750 nanograms of biotinylated and re-folded RNA (measured by NanoDrop) was incubated for 30 minutes shaking at 1.500 rpm at room temperature with protein extract (500 µg of whole cell protein extract or 150 µg of CPE) and murine RNase Inhibitor (NEB) in the corresponding protein extraction buffer (without glycerol). Streptavidin beads were added to the RNA pull down reaction mixture, followed by incubation for 30 minutes shaking at 1.500 rpm at room temperature. Afterwards, streptavidin beads were washed three times with ice-cold protein extraction buffer (whole cell or cytoplasmic, without glycerol) for five minutes each while rotating at room temperature and twice with 1x PBS for five minutes rotating at room temperature. The beads were eluted by incubation for three minutes at 95°C with SDS-sample buffer.

### Proteolytic in-gel digest

The eluates from 5’ tsRNA-Glu^CUC/UUC^ protein pull-down experiments were loaded on a SDS-PAGE and briefly stained with Coomassie Brilliant Blue. Stacked proteins were excised and gel pieces were washed, disulfide bridges were reduced with dithiothreitol and free thiols were alkylated with iodoacetamide. Proteins were digested overnight with trypsin (Promega) at 37°C. After digestion, peptides were extracted from the gel and cleaned-up on custom-made C18-stage tips^69^.

### Liquid chromatography – mass spectrometry (LC-MS/MS) for peptide identification

Tryptic digests were separated on an Ultimate 3000 RSLC nano-flow chromatography system (ThermoFisher Scientific), using a pre-column for sample loading (PepMapAcclaim) and a C18, 2 cm x 0.1 mm, 5 μm) and a C18 analytical column (PepMapAcclaim C18, 50 cm x 075 mm, 2 μm, Dionex-Thermo-Fisher Scientific), applying a linear gradient from 2 to 35 % solvent B (80 % acetonitrile, 0.1 % formic acid; solvent A 0.1 % formic acid) at a flow rate of 230 nL/min over 60 minutes. Eluting peptides were analyzed on a Q-Exactive HFX Orbitrap mass spectrometer, equipped with a Proxeon nanospray source (all ThermoFisher Scientific). For the data-dependent mode survey scans were obtained in a mass range of 375–1.500 m/z with lock mass on, at a resolution of 60.000 at 200 m/z and an AGC target value of 3E6. The 8 most intense ions were selected with an isolation width of 1.6 Da, fragmented in the HCD cell at 28 % collision energy and the spectra recorded at a target value of 1E5 and a resolution of 30.000. Peptides with a charge of +1 were excluded from fragmentation, the peptide match and exclude isotope features were enabled and selected precursors were dynamically excluded from repeated sampling for 15 seconds.

### Data analysis of protein identification

Raw data were searched with MaxQuant software package 1.6.0.16^70^ against UniProt human reference database (proteome ID: UP000005640) and a costum contaminant database with tryptic specificity allowing two missed cleavages. Carbamidomethylation was set as fixed modification, oxidation of methionine and N-terminal protein acetylation as variable modifications. All other parameters were set to default. LFQ feature and match between runs were activated. Results were filtered at a protein and peptide false discovery rate of 1 % at PSM and protein level. In Perseus 1.6.2.1, decoys and contaminants were filtered out, and the protein list reduced to entries with a minimum of one unique and razor peptide and a minimum of three LFQ values over 12 LC-MS/MS runs. LFQ values were log_2_-transformed, missing quantification values were replaced by a fixed value of 20. The protein intensity ratios (log_2_) for endo versus noRNA or scrambled RNA controls were calculated within replicates and plotted per condition. For stress-control comparison, protein hits with a positive log-fold change in both replicates and a log-fold change ≧1 in at least one of the two replicates under stress conditions were selected and represented in heatmaps.

### Immuno-fluorescence experiments

HEK293T-ecANG cells were plated onto poly-lysine-coated coverslips and Dox-mediated ecANG expression was induced for 48 hours. Cells were briefly rinsed in ice-cold PBS, fixed with 4 % PFA/1x PBS, washed with 1x PBS, blocked in blocking solution (3 % bovine serum albumin, 0.1 % Tx-100, 1x PBS) and stained with antibodies against human ANG in a wet chamber for 12 hours. Cells were washed in blocking solution followed by incubation with secondary goat-anti-mouse Alexa488-coupled secondary antibodies (1:500) for two hours at room temperature. Cells were washed in 1x PBS, exposed to Hoechst for DNA staining, embedded and imaged on a confocal microscope (Olympus FV3000).

### Determining tsRNA copy numbers in HEK293 cells

A mass dilution of both purified 5’ tsRNAs (measured using NanoDrop, ThermoScientific) was northern-blotted using probes against the 5’ halves of tRNA-Gly^GCC^ and tRNA-Glu^CUC^ along with total cellular RNA extracted from HEK293 cells growing under steady state conditions and from cells exposed to iAs followed by immediate harvesting and RNA extraction. In parallel, the mass of RNA after total RNA extraction using Trizol from 1*10^6^ HEK293 cells was determined in triplicates using NanoDrop. Radiographic signals from the dilution series of 5’ tsRNAs were quantified using *ImageJ* and plotted as a standard curve using Excel. The signals collected from HEK293 cells were subtracted (stress minus steady state) to arrive at a mass of tsRNAs after iAs-exposure. The calculated mass of tsRNAs was used to arrive at the number of tsRNA molecules normalized to the mass of loaded total RNA. The number of tsRNA molecules per cell was calculated using the equation: moles ssRNA (mol) = mass of ssRNA (g)/(length of ssRNA (nt)* 321.47 g/mol) + 18.02 g/mol) where the length for a tRNA half was set to 35 nt (the anticodon triplet being nucleotides 34 to 36). RNA copy number was calculated as moles of ssRNA* 6.022*e^23^ molecules/mol.

### Primers and Oligonucleotides (5’ to 3’)

ecANG construct primers: ANG_Gateway_fwd: AD0004:

GGGGACAAGTTTGTACAAAAAAGCAGGCTTCACCATGGTGATGGGCCTGGGCG

ANG_Gateway_rev: AD0005:

GGGGACCACTTTGTACAAGAAAGCTGGGTTCTAAGCGTAATCGGGAACATCG

*DNAzymes:*

V1: MATT_VIE361: AACCCGGGCCTCCCGCGTGG_GGCTAGCTACAACGA_AGGCGAGAATTCTACCACTG

V2: MATT_VIE362: TCGAACCCGGGCCTCCCGCG_GGCTAGCTACAACGA_GGCAGGCGAGAATTCTACCA

*in vitro transcription (IVT)/ribozyme single-stranded DNA template:*

scrambled small RNA: AD114: GGTGACTGGAGTTCAGACGTGTGCTCTTCCGATCTGACGGTACCGGGTACCGTTTCGTCCTCACGGACTCATCAGAGATCGGAATCTCCCTATAGTGAGTCGTATTA

*PCR primers for making double-stranded IVT/ribozyme templates:* scrambled small RNA_fwd: AD115: GGTGACTGGAGTTCAGACGTG universal_rev: MATT_VIE174: CGCGCGAAGCTTAATACGACTCACTATA

*Northern blotting probes:*

5’ tRNA-Gly^GCC^: AD0006: TCTACCACTGAACCACCAAT

3’ tRNA-Gly^GCC^: AD0007: TGGTGCATTGGCCGGG

5’ tRNA-Glu^CUC^: AD0008: GAATCCTAACCACTAGACCAC

*RNA affinity capture probes:*

tRNA-Glu^CUC^: AD0022: 5’-Amino-Modifier C6-CGCCGAATCCTAACCACTAGACCACCA tRNA-Gly^GCC^: AD0023: 5’-Amino-Modifier C6-AGGCGAGAATTCTACCACTGAACCACC

