## Supplementary material for "Production and Purification of Endogenously Modified tRNA-Derived Small RNAs": STable 1

| Compound | Precursor Ion | MS1 Res | Product Ion | MS2 Res | Ret Time [min] | Fragmentor [V] | Collision Energy [eV] |
|---|---|---|---|---|---|---|---|
| A | 268.1 | Wide | 136 | Unit | 5.1 | 200 | 25 |
| A SILIS | 283 | Wide | 146 | Unit | 5.1 | 200 | 25 |
| Am | 282.1 | Wide | 136 | Unit | 6.1 | 130 | 17 |
| Am SILIS | 298 | Wide | 146 | Unit | 6.1 | 130 | 17 |
| C | 244.1 | Wide | 112 | Unit | 2 | 200 | 20 |
| C SILIS | 256 | Wide | 119 | Unit | 2 | 200 | 20 |
| G | 284.1 | Wide | 152 | Unit | 4.1 | 200 | 25 |
| G SILIS | 299 | Wide | 162 | Unit | 4.1 | 200 | 25 |
| m1A | 282.1 | Wide | 150 | Unit | 2.3 | 150 | 25 |
| m1A SILIS | 298 | Wide | 161 | Unit | 2.3 | 150 | 25 |
| m1G | 298.1 | Wide | 166 | Unit | 4.8 | 105 | 13 |
| m1G SILIS | 314 | Wide | 177 | Unit | 4.8 | 105 | 13 |
| m22G | 312.1 | Wide | 180 | Unit | 5.6 | 105 | 13 |
| m22G SILIS | 329 | Wide | 192 | Unit | 5.6 | 105 | 13 |
| m2G | 298.1 | Wide | 166 | Unit | 5 | 95 | 17 |
| m2G SILIS | 314 | Wide | 177 | Unit | 5 | 95 | 17 |
| m5C | 258.1 | Wide | 126 | Unit | 3.5 | 185 | 13 |
| m5C SILIS | 271 | Wide | 134 | Unit | 3.5 | 185 | 13 |
| m5U | 259.1 | Wide | 127 | Unit | 4.2 | 95 | 9 |
| m5U SILIS | 271 | Wide | 134 | Unit | 4.2 | 95 | 9 |
| m6A | 282.1 | Wide | 150 | Unit | 6.5 | 125 | 17 |
| m6A SILIS | 298 | Wide | 161 | Unit | 6.5 | 125 | 17 |
| m7G | 298.1 | Wide | 166 | Unit | 3.3 | 100 | 13 |
| m7G SILIS | 314 | Wide | 177 | Unit | 3.3 | 100 | 13 |
| U | 245.1 | Wide | 113 | Unit | 2.9 | 95 | 5 |
| U SILIS | 256 | Wide | 119 | Unit | 2.9 | 95 | 5 |
| Um | 259.2 | Wide | 113 | Unit | 4.4 | 96 | 8 |
| Um SILIS | 271.1 | Wide | 119 | Unit | 4.4 | 96 | 8 |
| Y | 245.1 | Wide | 209 | Unit | 1.7 | 90 | 5 |
| Y SILIS | 256 | Wide | 220 | Unit | 1.7 | 90 | 5 |
