## Supplementary material for "Production and Purification of Endogenously Modified tRNA-Derived Small RNAs": STable 2

**iAs**

| | tRNA Gly-1 | tRNA Gly-2 | tRNA Gly-3 | tsRNA Gly-1 | tsRNA Gly-2 | tsRNA Gly-3 | tRNA Glu-1 | tRNA Glu-2 | tRNA Glu-3 | tsRNA Glu-1 | tsRNA Glu-2 | tsRNA Glu-3 |
|---|---|---|---|---|---|---|---|---|---|---|---|---|
| **Um** | 0,29560801 | | 0,28684211 | 0,41349737 | 0,54161162 | 0,31247111 | | 0,1068281 | 0,11878771 | 0,00048628 | 0,00332907 | 0,00140809 |
| **m2G** | 0,54814731 | 0,87347984 | 0,41842105 | 0,58042779 | 0,51005431 | 0,46731996 | 0,60874258 | 0,44707558 | 0,37751712 | 0,63435821 | 0,50564934 | 0,49651424 |
| **Psi** | 0,4959428 | 0,67719223 | 0,64605263 | 0,020292 | 0,14604433 | 0,07303319 | 0,81079331 | 0,96946497 | 0,95572581 | 0,48433943 | 0,43080682 | 0,44203198 |
| **m1A** | 0,37587244 | 0,28952422 | 0,30328947 | 0,00484966 | 0,02421841 | 0,01941389 | 0,20205073 | 0,17786878 | 0,16814699 | 0,00972569 | 0,00480866 | 0,00639056 |
| **m5C** | 1,66467117 | 0,78515042 | 1,19605263 | 0,01250702 | 0,03522677 | 0,03512989 | 0,73826228 | 1,09338556 | 0,82554749 | 0,01774939 | 0,00838433 | 0,01158966 |
| **m5U** | 0,3151847 | 0,16684446 | 0,30131579 | 0,00063811 | 0,00660502 | 0,00138671 | 0,01813276 | 0,03258257 | 0,04881687 | 0,00291771 | 0,0001233 | 0,00303281 |
| **m7G** | 0,02871248 | 0,13740132 | 0,04078947 | 0,00370106 | 0,02054895 | 0,0143293 | 0,09325418 | 0,05394819 | 0,06780121 | 0,00753741 | 0,00369897 | 0,00498247 |
| **m1G** | | | | 0,001 | | | | | 0,0012 | 0,0442 | 0,0787 | 0,1645 |
| **m22G** | 0,0358906 | 0,20119479 | 0,04802632 | 0,00523253 | 0,03008953 | 0,02033836 | 0,1295197 | 0,06623342 | 0,08678554 | 0,01069826 | 0,00493196 | 0,00530742 |
| **Am** | 0,02479714 | 0,18647322 | 0,04407895 | 0,00459442 | 0,02788786 | 0,02449847 | 0,09066379 | 0,0197632 | 0,04339277 | 0,01045512 | 0,00456206 | 0,00758202 |
| **m6A** | 0,04828917 | 0,03925752 | 0,05065789 | 0,00063811 | 0,0080728 | 0,00231118 | 0,03367512 | 0,02830945 | 0,03145976 | 0,00121571 | 0,0006165 | 0,00097483 |

**ecANG**

| | tRNA Gly-1 | tRNA Gly-2 | tRNA Gly-3 | tsRNA Gly-1 | tsRNA Gly-2 | tsRNA Gly-3 | tRNA Glu-1 | tRNA Glu-2 | tRNA Glu-3 | tsRNA Glu-1 | tsRNA Glu-2 | tsRNA Glu-3 |
|---|---|---|---|---|---|---|---|---|---|---|---|---|
| **Um** | 0,14787485 | 0,13453238 | 0,18572594 | 0,19811395 | 0,05574847 | 0,30295211 | 0,12037109 | 0,0665521 | 0,93440318 | | | |
| **m2G** | 0,35283123 | 0,52931485 | 0,67291641 | 0,89636868 | 0,29171024 | 1,2234365 | 0,83607194 | 0,66226488 | 0,58040767 | 0,80740652 | 0,72363281 | 1,07104837 |
| **Psi** | | | 1,42515643 | 0,13962744 | 0,36419559 | 0,05162786 | | | 2,29120426 | 0,65226147 | 0,38631076 | 1,07697828 |
| **m1A** | 0,25254462 | 0,30410553 | 0,39862956 | 0,00784191 | 0,05598071 | 0,00613558 | 0,3055795 | 0,27160965 | 0,24115019 | 0,01818351 | 0,00971101 | 0,02118258 |
| **m5C** | 0,90021374 | 1,03012182 | 0,91225084 | 0,03156662 | 0,05635556 | 0,02011205 | 0,50360295 | 0,56540333 | 0,69290973 | 0,03723759 | 0,03940481 | 0,07041207 |
| **m5U** | 0,23130119 | 0,31284757 | 0,25265908 | 0,02354001 | 0,04791899 | 0,00439129 | 0,0231189 | 0,071492 | 0,0683735 | 0,02343906 | 0,01984596 | 0,01915426 |
| **m7G** | 0,07259732 | 0,0837392 | 0,18202531 | 0,00243188 | 0,01842332 | 0,00208065 | 0,15756443 | 0,15048643 | 0,10335008 | 0,00650504 | 0,00430749 | 0,01107916 |
| **m1G** | 0,11524738 | 0,01434196 | 0,4957748 | | | | 0,03141267 | 0,03686564 | 0,01729083 | | | |
| **m22G** | 0,07278896 | 0,01762931 | 0,27039675 | 0,01020171 | 0,02600582 | 0,00226228 | 0,03342276 | 0,02936321 | 0,07029088 | 0,0086519 | 0,01468888 | 0,02442693 |
| **Am** | 0,05540978 | 0,09035019 | 0,00663629 | 0,01275039 | 0,04296532 | 0,00374186 | 0,18395092 | 0,14681603 | 0,10056844 | 0,00839067 | 0,02781475 | 0,0259476 |
| **m6A** | 0,0351445 | 0,02315662 | 0,06682964 | 0,00638078 | 0,01208399 | 0,00142595 | 0,01432404 | 0,0146816 | 0,02120002 | 0,00499797 | 0,00873754 | 0,01236703 |
